# Characterisation of the transcriptional dynamics underpinning the function, fate, and migration of the mouse Anterior Visceral Endoderm

**DOI:** 10.1101/2021.06.25.449902

**Authors:** Shifaan Thowfeequ, Jonathan Fiorentino, Di Hu, Maria Solovey, Sharon Ruane, Maria Whitehead, Bart Vanhaesebroeck, Antonio Scialdone, Shankar Srinivas

## Abstract

During early post-implantation development of the mouse embryo, the Anterior Visceral Endoderm (AVE) differs from surrounding visceral endoderm (VE) in its migratory behaviour and ability to restrict primitive streak formation to the opposite side of the egg cylinder. In order to characterise the molecular basis for the unique properties of the AVE, we combined single-cell RNA-sequencing of the VE prior to and during AVE migration, with high-resolution imaging, short-term lineage labelling, phosphoproteomics and pharmacological intervention. This revealed the transient nature of the AVE, the emergence of heterogeneities in AVE transcriptional states relative to position of cells, and its prominence in establishing gene expression asymmetries within the spatial constraints of the embryo. We identified a previously unknown requirement of Ephrin- and Semaphorin-signalling for AVE migration. These findings point to a tight coupling of transcriptional state and position in the AVE and reveal molecular heterogeneities underpinning its migratory behaviour and function.

## INTRODUCTION

Axial specification is a relatively late event during mammalian embryogenesis, as extraembryonic tissues, essential for this process, need to be established first. One such tissue, known to play a pivotal role in the specification of anterior–posterior pattern, is the visceral endoderm (VE). The mouse VE, equivalent to the hypoblast in other amniotes, is a simple epithelium that encapsulates both the pluripotent epiblast and the extra-embryonic ectoderm (ExE) (Gardner and Rossant, 1979; Hermitte and Chazaud, 2014; Stern and Downs, 2012). The VE plays both a trophic and an instructive role, by nourishing and patterning the underlying tissues respectively (Bielinska et al., 1999). Around Embryonic Day (E) 5.5, a morphologically distinct sub-region of the embryonic VE, termed the anterior visceral endoderm (AVE; also called the distal visceral endoderm at this stage based on its initial position at the distal tip of the egg cylinder) is specified (Lawson and Wilson, 2016; Stower and Srinivas, 2018; Thomas and Beddington, 1996). In response to unknown cues, the AVE initiates a characteristic unidirectional migration towards the presumptive anterior region of the embryo, and over a period of 4-5 hours reaches the boundary between the epiblast and ExE (Srinivas et al., 2004). From this position, AVE cells secrete inhibitors of the Wnt and Nodal signalling pathways (DKK1, CER1, and LEFTY1), thereby restricting primitive streak (PS) formation to the opposite side of the egg cylinder (Argelaguet et al., 2019; Brennan et al., 2001; Kimura et al., 2000; Kimura-Yoshida et al., 2005; Yamamoto et al., 2004) and establishing the earliest features of embryonic anterior– posterior (A–P) polarity. Cells from the distal tip of the egg cylinder continue to migrate to the presumptive anterior so that by E6.25, the original AVE cells to occupy the anterior position have been displaced laterally and replaced by later arriving AVE cells (Migeotte et al., 2010; Srinivas et al., 2004; Takaoka et al., 2011).

AVE cells express characteristic markers of ‘anterior’ pattern while still at the distal tip of the egg cylinder. The correct directional migration of the AVE from the distal tip of the egg cylinder to the boundary with the ExE, resolves an ‘A–P’ axis (as defined by marker expression) that is initially aligned with the proximal–distal (P–D) axis of the egg cylinder, to a definitive one that is orthogonal to it (Arnold and Robertson, 2009). Mutants in which these cells fail to migrate, or do so improperly, show defects in the localised formation of the PS (Stower and Srinivas, 2018). A subset of the human hypoblast that is equivalent to the mouse AVE has recently been discovered indicating a conservation of the function of the AVE across mammals (Molè et al., 2021). AVE cells show characteristics of active migration such as polarized cellular projections (Srinivas et al., 2004). However, the entire VE remains an intact simple epithelial monolayer during their migration (Trichas et al., 2011), so AVE cells must negotiate their way through the surrounding VE cells to assume their anterior position. The actomyosin cytoskeleton plays a central role during this characteristic migration (Bloomekatz et al., 2012; Migeotte et al., 2010; Omelchenko et al., 2014; Rakeman and Anderson, 2006). The extracellular and intercellular cues controlling AVE migration are much less clear. For example, exogenously supplied DKK1 can act as a guidance cue in this process (Kimura-Yoshida et al., 2005), however the precise role of endogenous DKK1 is unclear given that it is not required for AVE migration (Mukhopadhyay et al., 2001).

Though the VE is a single continuous sheet of broadly similar cells, there is evidence that there might be underlying heterogeneity amongst them. Cells in the VE overlying the epiblast (referred to as ‘embryonic’ VE or emVE) show significantly greater dynamic behaviour and morphological irregularity in cell shapes than the VE cells overlying the extraembryonic ectoderm (exVE). AVE cells, from the time of their induction, are also morphologically different from surrounding emVE cells, being distinctly columnar (Rivera-Pérez et al., 2003; Srinivas et al., 2004). However, it is not known whether the initial population of AVE cells, once they become displaced laterally, remain distinct from surrounding VE cells and, for example, retain their capacity to restrict PS formation. AVE cells are also different from neighbouring emVE cells in expressing a specific repertoire of markers (such as *Hhex*, *Hesx1*, *Lhx1*, *Otx2*, *Lefty1* and *Cer1* (Hoshino et al., 2015; Stower and Srinivas, 2018). A systematic and unbiased characterization of the underlying transcriptional profiles that underpin the unique properties of AVE cells, and the order in which they emerge during anterior patterning, is thus far lacking.

We therefore combined full-length, high-coverage single-cell RNA-sequencing with high-resolution imaging of the VE prior to and during AVE migration to characterize differences between AVE cells and surrounding emVE cells in order to: (1) determine how the molecular properties of AVE cells change as they migrate directionally to establish the A–P axis and pattern the underlying epiblast; (2) identify new signalling pathways that regulate this unique migratory behaviour in the early post-implantation embryo.

We identified a previously unappreciated degree of transcriptional heterogeneity within the VE across both space and the relatively short developmental timescales of AVE migration. We characterized the transient transcriptional state of AVE cells during their emergence and migration, established their ‘fate’ once they had completed migration, and validated these findings using lineage labelling. Using a combination of multiplexed-Hybridization Chain Reaction and high-resolution imaging, we found that polarized expression of markers in the prospective anterior VE precedes that in the posterior VE and the epiblast. Finally, we identified signalling pathways likely to play important roles during anterior patterning and experimentally verified the previously unknown requirement of two pathways—Ephrin and Semaphorin signalling—for AVE migration.

## RESULTS

### Single-cell transcriptomic profiling and proteomic analyses identify novel markers of visceral endoderm (VE) subpopulations

We collected single cells from two stages of mouse embryos, E5.5 and E6.25, that bookend the start and end of AVE migration, respectively. To obtain a VE-enriched collection of single cells, prior to disaggregation, we labelled embryos with a fluorescent membrane dye that was taken up primarily by the outer cell layers of the embryo. As embryos at these stages have fewer than 200 to 250 VE cells each, we pooled up to 10 stage-matched embryos for each collection. Labelled cells from disaggregated embryos were isolated using FACS, and processed through the Smart-seq2 protocol, which allows full-length, high-coverage single-cell RNA-sequencing (scRNA-seq) (Picelli et al., 2013) (Figure 1A). After quality control, 252 cells at E5.5 and 235 at E6.25 were retained for downstream analysis. We performed unsupervised clustering of the cells (see Methods), which led to the identification of five clusters per stage (Figure 1B). On the basis of the expression pattern of known marker genes, the two smallest clusters corresponded to epiblast (Epi) and extra-embryonic ectoderm (ExE) cells (Figure 1C). The remaining three larger clusters at each stage represented sub-types of the VE (positive for *Gata6* and *Amn*). Using known combinations of marker genes, we further annotated them as early- or late-AVE (at E5.5 and E6.25 respectively, both expressing high levels of *Cer1*, *Lefty1*, and *Hhex*), emVE (i.e., the non-migratory VE cells overlying the epiblast, with lower expression of *Cer1, Lefty1*, and *Hhex*), and exVE (i.e., the VE cells overlying the ExE, with the highest expression of *Cubn* and low expression of *Afp*; Figure 1C, Supplementary Figure 1A). These results indicated that for both stages, VE-enrichment was successful, as the majority of cells were associated with the VE clusters while Epi and ExE cells formed the two smaller clusters. Additionally, we identified marker genes of each cluster in an unbiased way, which revealed new markers particularly for the different VE clusters (Supplementary Figure 1B, Supplementary Tables 1 and 2).

**Figure 1.**
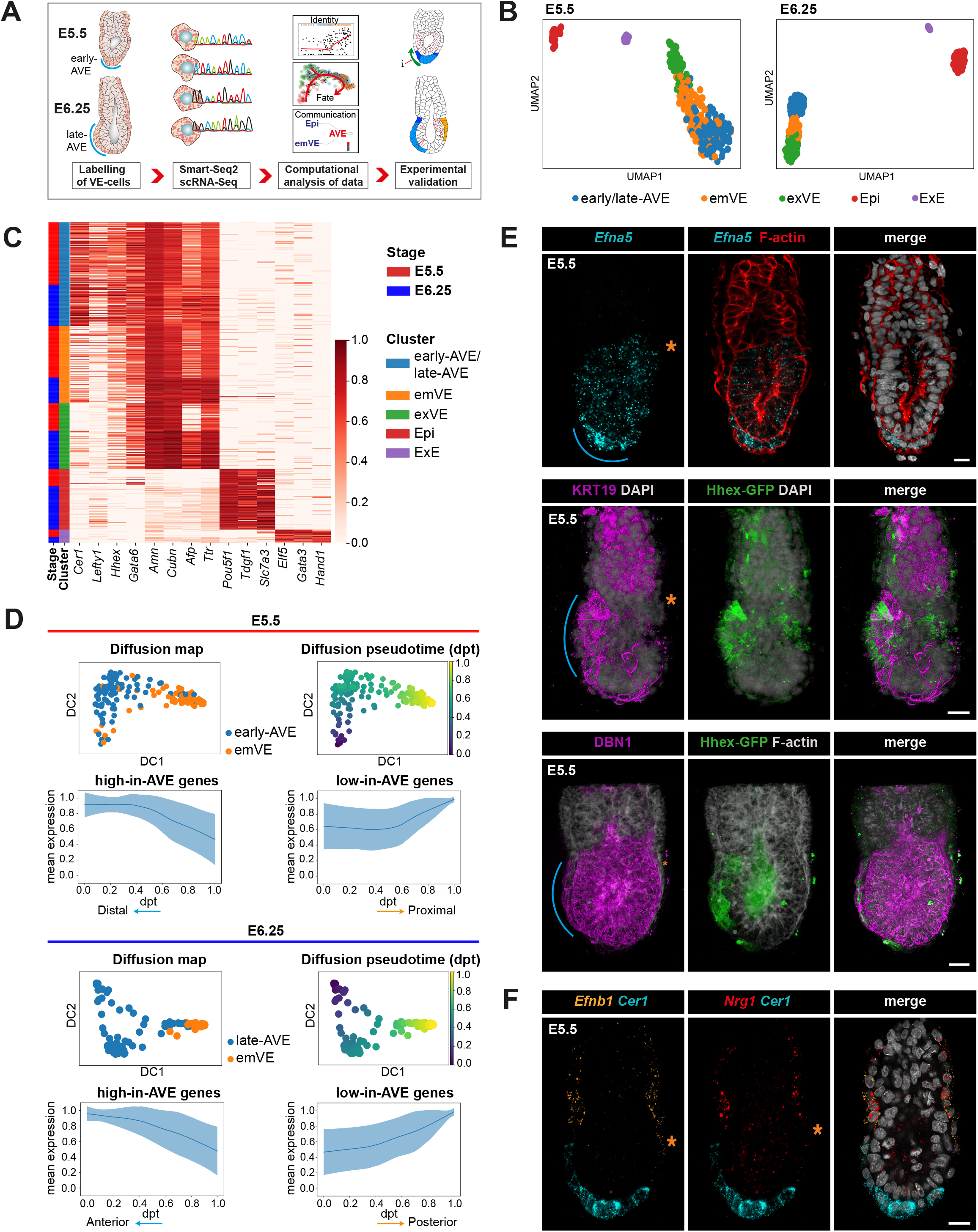
Computational analysis of a visceral endoderm-enriched single-cell RNA-seq dataset of mouse embryos at embryonic days 5.5 and 6.25 and experimental identification of novel anterior and posterior VE markers. (A) Schematic summarising this study for the enriched isolation of VE cells for Smart-seq2, post-sequencing computational analyses and experimental validation of the results. (B) UMAP plots of the cells that passed quality control from E5.5 (n=252) and E6.25 (n=235) embryos. Cells were clustered in the same five groups at each stage, as indicated by the different colours: early- or late-anterior visceral endoderm (AVE), rest of the VE cells surrounding the epiblast and ExE (emVE and exVE respectively), the epiblast (Epi), the extraembryonic ectoderm (ExE). (C) Heatmap showing the expression of known marker genes for the identified cell types at E5.5 and E6.25, with the cells grouped according to the clustering shown in (B). The normalized log expression levels for each gene are standardised so that they vary within the interval [0,1]. (D) First two diffusion components (DC1 and DC2) of a diffusion map computed for early- or late-AVE and emVE cells, coloured according to the cluster (left) and to a diffusion pseudotime (dpt) coordinate (right), and mean standardised expression of the genes belonging to ‘high-in-AVE’ and ‘low-in-AVE’ groups, at E5.5 (top) and E6.25 (bottom). The gene clusters are obtained through a hierarchical clustering of the genes that show a significant trend in diffusion pseudotime, selected using a Generalised Additive Model. The dpt axis tracts the distal–proximal (at E5.5) or anterior–posterior (at E6.25) axis of the embryo in the directions indicated by the arrows. (E) Validation of the expression of newly identified ‘high-in-AVE’ markers using *in situ* HCR (for *Efna5*) or immunofluorescence (for KRT19 and DBN1). Hhex-GFP marks the AVE. The blue line indicates the position of the AVE and the orange asterisk marks the posterior side of the embryo. (F) Validation of the expression of newly identified low-in-AVE markers (*Efnb1* and *Nrg1*) at E5.5 using *in situ* HCR. *Cer1* marks the AVE and the orange asterisk marks the posterior side of the embryo. All scale bars represent 20 *μ*m and all embryos are orientated with the anterior on the left (←).

To understand how the transcriptional profiles of the different VE sub-types related to each other, we analysed the relative distance and the connectivity between the VE clusters with the Partition-based graph abstraction (PAGA) algorithm (Wolf et al., 2019). This showed that the VE clusters were highly connected, with the AVE at both stages being most closely connected to the respective emVE at that stage (Supplementary Figure 1C, D). The analysis also indicated that the AVE at E6.25 is transcriptionally more distinct from the rest of the VE than at E5.5.

We then computed separate diffusion maps (Haghverdi et al., 2015) at the two stages, to identify differences between the transcriptomes of AVE and emVE cells at E5.5 and E6.25, given the subtlety in their distinction. Cells were ordered through the definition of a diffusion pseudotime (dpt) coordinate (Haghverdi et al., 2016) originating from the respective AVE cluster at each stage (Figure 1D). We identified the genes differentially expressed in pseudotime, and using an unsupervised clustering analysis (see Methods), split them into two gene groups based on their high or low expression in the AVE cell clusters (‘high-in-AVE’ and ‘low-in-AVE’ gene groups; Figure 1D, Supplementary Tables 3 and 4). The presence of well-known AVE markers among the ‘high-in-AVE’ genes (e.g., *Cer1*, *Hhex*) suggests that the pseudotime axis (dpt) appears be tracking the spatial position of cells along the A–P axis at E6.25, and the P–D axis at E5.5.

To test this hypothesis, we chose a number of novel marker genes from the ‘high-in-AVE’ and ‘low-in-AVE’ groups common to both stages, to determine their spatial expression patterns in E5.5 and E6.5 embryos via multiplexed-*in situ* Hybridisation Chain Reaction (HCR) and wholemount immunofluorescence (Figure 1E and F, Supplementary Figure 1B, E, and 2A-C). In these assays, AVE cells were independently identified based either on their distinct columnar morphology or by the expression of reference AVE markers such as *Cer1* or the Hhex-GFP transgene (Rodriguez et al., 2001). *Efna5*, *Krt19* and *Dbn1*, which are in the ‘high-in-AVE’ group at both stages were chosen for further validation based on their high specificity to the AVE clusters, novelty as AVE-markers and as they code for proteins with potential links to cell migration. *Efna5* transcript, as well as KRT19 and DBN1 proteins were all expressed at high levels in AVE cells at E5.5 (Figure 1E) and E6.25 (Supplementary Figure 2A–C) with little expression in surrounding emVE cells. Therefore, these stains confirmed that the dpt coordinate did indeed represent the P–D and A–P axes of the E5.5 and E6.25 embryo respectively. Additionally, KRT19 and DBN1 were also expressed in the ExE and the epiblast respectively, in agreement with the scRNA-seq dataset (Figure 1E, Supplementary Figure 1B, 2B and C).

Next, we validated two genes in the ‘low-in-AVE’ groups at both stages, *Nrg1* and *Efnb1* (Figure 1F, 5H, and Supplementary Figure 1E). *Nrg1* and *Efnb1* were both highest in the exVE (as also observed in our scRNA-seq data, see Supplementary Figure 1B), and their expression domains extended distally into the emVE only on the presumptive-posterior, the side opposite to the AVE at E5.5 and E6.5 (Figure 1F and 5H). These results further corroborate the interpretation of the diffusion pseudotime axis as a spatial axis, spanning the P–D and A–P axes at E5.5 and E6.25 respectively.

Transcriptomic differences have functional significance largely only in so far as they are translated to the proteome. Hence, to determine the extent to which the transcriptional profile of a cell might reflect its functional protein complement, we compared our single-cell transcriptomic data with mass spectrometry-driven proteomic and phosphoproteomic data from E6.5 embryos. We isolated E6.5 embryos and, in order to retain some spatial information, bisected them along the epiblast-ExE boundary into ‘embryonic’ (epiblast and overlying emVE) and ‘abembryonic’ (ExE and overlying exVE) portions (see Supplementary Figure 1A) before separately profiling the proteome and phosphoproteome of these two regions. After a preliminary filtering of the peptides, aggregation to the protein level and sample normalisation, differentially expressed proteins between the embryonic and abembryonic halves were identified (see Methods; Supplementary Figure 2D–F and Supplementary Tables 5-8).

A comparison showed that overall, there was good agreement between the proteomic and the transcriptomic data, as shown by the statistically significant correlation between the log2-fold changes computed from the two datasets (see Methods and Supplementary Figure 2E). However, we also identified some proteins such as KMT2A and PML that showed differences in phosphorylation but were transcriptionally expressed uniformly across the VE (Supplementary Figure 2F), illustrating the utility of the phosphoproteomic data set in complementing our scRNA-Seq data for the identification of candidates important in AVE migration and function. Interestingly, the data also revealed that, among the ‘high-in-AVE’ genes, there are some associated with cell migration, such as *Dbn1*, *Marcks*, *Marcksl1*, and *Stmn1* (El Amri et al., 2018; Ni et al., 2017; Tanabe et al., 2014; see Supplementary Figure 2B and F), whose associated protein show an upregulation in the phosphoproteomic data from the embryonic halves, suggesting that they may also be subject to post-translational regulations.

### Order of emergence of asymmetric gene expression along the A–P axis

Our diffusion analysis captures the transformation of a P–D asymmetry in marker expression into an A–P asymmetry in expression. We therefore used it to investigate the sequence in which asymmetric gene expression emerges in the VE. It has been suggested that even before the anterior shift in AVE markers resulting from the migration of cells expressing them, *Wnt3* is expressed exclusively in the posterior VE in a localised manner (Rivera-Pérez and Magnuson, 2005). To test this possibility, we first looked at our diffusion analyses. These showed that at both E5.5 and E6.25, in comparison to the posterior specific asymmetric expression of *Wnt3*, the anterior specific asymmetric expression of *Cer1* is more pronounced and robust (Figure 2A). To establish the relative timing of the asymmetric expression of these two genes, we used multiplexed-HCR to detect both *Cer1* (distal/anterior) and *Wnt3* (proximal/posterior) simultaneously in the same embryos. We found that in all E5.5 embryos where *Cer1* was expressed symmetrically at the distal tip, *Wnt3* was also expressed symmetrically across the proximal egg cylinder (both epiblast and emVE) (N=6; Figure 2B, ‘pre-migration’). In embryos where *Cer1* had started to show clearly asymmetric localisation to the prospective anterior as a result of AVE migration, *Wnt3* expression still spanned the proximal portion of the egg cylinder, and was not just isolated to the posterior half (N=16; Figure 2B, ‘mid-migration’). We only observed a complete segregation of the *Wnt3* expression domain to the presumptive posterior in embryos staged beyond E6.0, before the PS had emerged, but after the *Cer1*-expressing AVE cells had reached the epiblast–ExE boundary in the same embryos (N=27; Figure 2B, ‘post-migration’). This indicates that axial pattern emerges first by the asymmetric expression of anterior markers within the VE, followed subsequently by asymmetric expression of posterior markers within the VE and epiblast.

**Figure 2.**
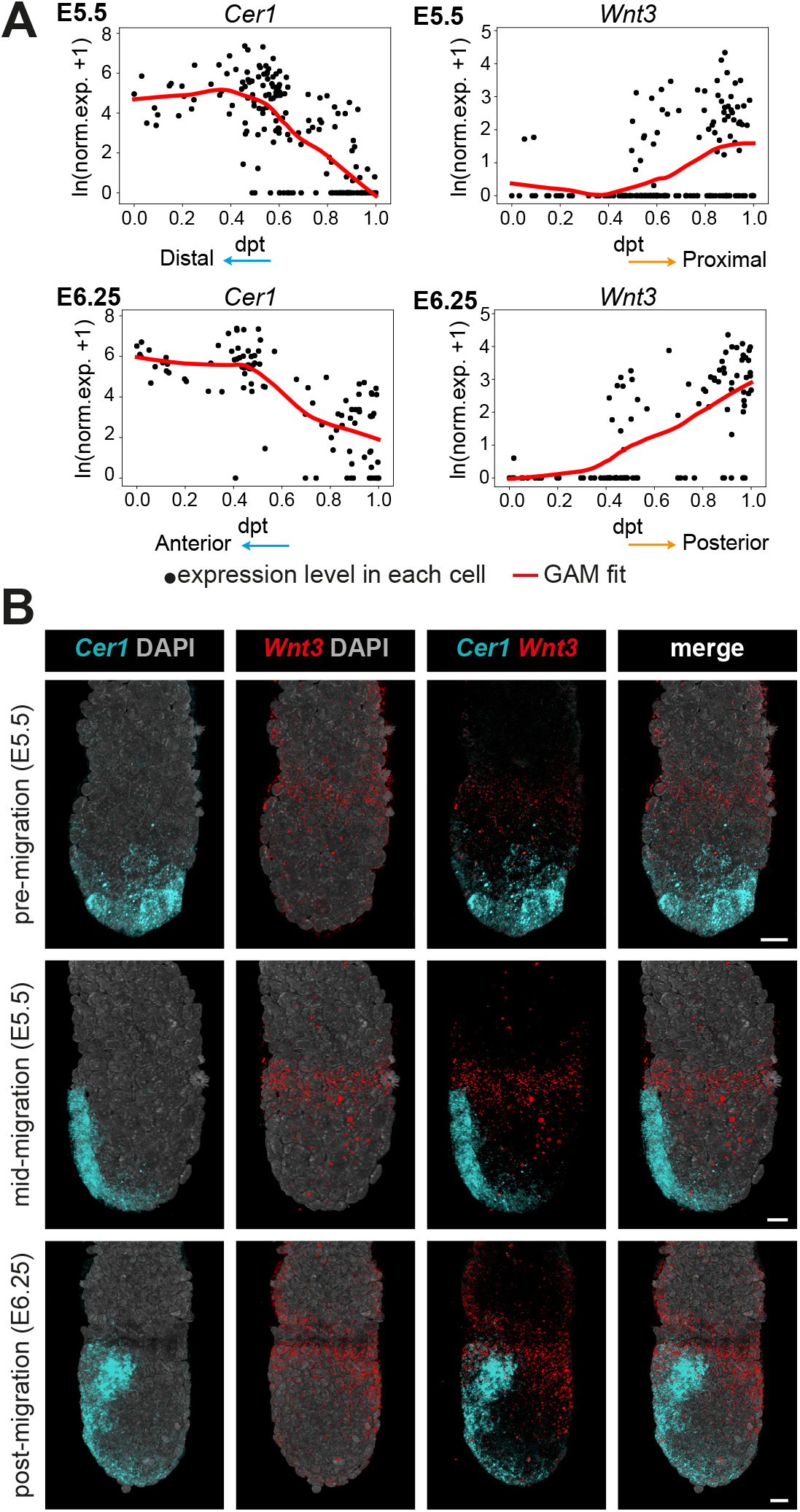
Symmetry breaking along the anterior–posterior axis relative to *Wnt3* expression in direct comparison to the migration of the AVE. (A) Expression patterns of *Cer1* and *Wnt3* in the AVE and emVE clusters, as a function of diffusion pseudotime, at E5.5 (top) and E6.25 (bottom). Each black dot represents a cell, the red line shows the fit obtained from a Generalised Additive Model (GAM). (B) Volume renderings showing the changes to the *Wnt3* expression domain (visualised by *in situ* HCR) relative to the position of the *Cer1*-expressing AVE cells as they migrate from the distal portion of the embryo to the epiblast–ExE boundary and then laterally across the embryo. Scale bars represent 20 *μ*m and all embryos are orientated with the anterior on the left (←).

Next, to test if DKK1 might act as a guidance cue for AVE cells (Kimura-Yoshida et al., 2005), we again looked at our diffusion pseudotime analyses and hypothesised that as a guidance cue, *Dkk1* expression should show clear anterior asymmetry prior to the asymmetric expression of canonical AVE markers such as *Cer1*. The diffusion pseudotime analyses showed that while *Cer1* marked the distal and anterior regions of VE before and after migration respectively (at E5.5 and E6.25), *Dkk1* became elevated in the anterior, only at E6.25 (Supplementary Figure 3A). To validate this result, we used HCR to visualize the emergence of the relative asymmetry in expression of *Dkk1* and of the AVE marker *Cer1*, again, within the same embryos. At E5.5, prior to AVE migration, *Dkk1* was expressed in cells interspersed throughout the distal tip and in a radially symmetrical ring proximal to the *Cer1*-expression domain (N=9). During AVE migration, *Dkk1* expression was restricted to two distinct domains in the anterior and posterior (N=10; Supplementary Figure 3B). It was only around E6.25, at later stages of AVE migration, that *Dkk1* expression became specifically enriched in the anterior (N=4), in agreement with our diffusion pseudotime analyses (Supplementary Figure 3A) and previous reports on the expression of DKK1 protein (Hoshino et al., 2015).

### Asymmetry in the VE emerges approximately 12–18 hours before it does in the epiblast

Axial asymmetry established in the VE by directional AVE migration is transferred to the epiblast by the AVE inhibiting the expression of PS-specific genes in the subjacent epiblast, thereby restricting PS formation to the opposite side of the epiblast. To determine the precise sequence of asymmetric marker expression in the VE and the underlying epiblast, we analysed the temporal expression patterns of the posterior marker *Nodal* in direct relation to the AVE marker *Cer1*. We used *Brachyury* (*T*) as a reference marker for the PS. Our single-cell data showed that in the epiblast, *Nodal* is expressed at both E5.5 and E6.25; at similar stages however, *T* is detected in only a very limited number of epiblast cells (Figure 3A and Supplementary Figure 4A). This indicates that *T* expression in the epiblast is induced well after AVE migration is already underway, consistent with it becoming robustly expressed only later, during primitive streak formation at ~E6.5 (Mohammed et al., 2017; Scialdone et al., 2016). In addition to expression in the epiblast, *Nodal* is also expressed in the VE, where it is important for the initial induction of the AVE (Tremblay et al., 2000). Our analyses unexpectedly revealed that at E6.25, *Nodal* was statistically significantly depleted in the posterior VE compared to the AVE (Figure 3A). We did not detect expression of *T* in any of the VE populations at any stage (Supplementary Figure 4A and B).

**Figure 3.**
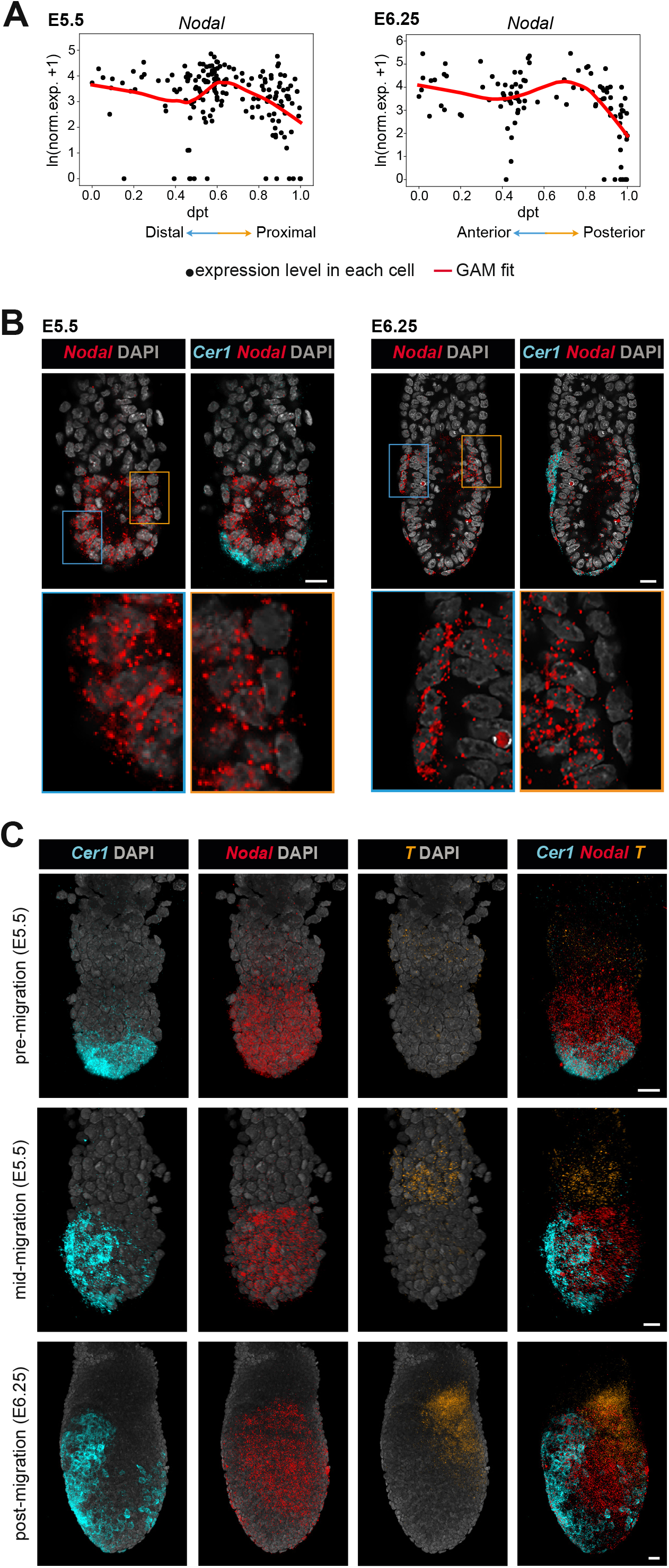
Symmetry breaking along the anterior–posterior axis within the epiblast and VE relative to *Nodal* and *T* expression in direct comparison to the migration of the AVE. (A) Expression patterns of *Nodal* in the AVE and emVE clusters, as a function of diffusion pseudotime, at E5.5 (left) and E6.25 (right). (B) Optical sections through embryos showing the distinct expression of *Nodal* in the epiblast and the VE relative to that of *Cer1* (visualised by *in situ* HCR) at E5.5 and E6.25. Individual magnifications of the anterior and posterior are shows underneath to highlight the distinct *Nodal* expression in the VE and epiblast within these regions. (C) Volume renderings showing the change in *Nodal* and *T* expression (visualised by *in situ* HCR) relative to the position of the *Cer1*-expressing AVE cells as they migrate from the distal portion of the embryo to the epiblast-ExE boundary and then laterally across the embryo. All scale bars represent 20 *μ*m and all embryos are orientated with the anterior on the left (←).

To validate these results, we performed multiplexed-HCR for *Nodal*, *Cer1* and *T* on embryos between E5.5 and E6.25. This verified that at both E5.5 and E6.25, *Nodal* expression in the VE is depleted specifically in the posterior VE, while showing high expression in the *Cer1*-positive AVE cells (N=6 and =4; Figure 3B). *Nodal* was detected uniformly throughout the epiblast as expected at E5.5, in embryos where *Cer1* was confined to the distal region (N=3; Figure 3C) or already slightly shifted towards the anterior (N=6; Figure 3B). *T* could not be detected in either the ExE or the epiblast at this stage (Figure 3C). At mid-migration stages, when *Cer1* was detected asymmetrically in the future anterior, *Nodal* started to show a very slight imbalance of expression towards the posterior epiblast, and *T* was detectable only in the ExE as previously reported (Rivera-Pérez and Magnuson, 2005), where it was expressed symmetrically (N=5). It was only at E6.25, when *Cer1*-expressing cells were already positioned along the anterior and lateral sides of the embryo, that *Nodal* started to be clearly restricted to the posterior of the epiblast (N=3). By this stage, *T* could also be detected in the epiblast, asymmetrically in a proximal posterior domain, as well in the posterior ExE (Figure 3C). These diffusion analyses and their subsequent HCR validation establish a sequence where robust asymmetry of marker expression in the epiblast only emerges at E6.25, approximately 12 to 18 hours after such asymmetry is seen in the VE at E5.5 as a result of AVE migration.

### Transcriptional dynamics indicates the origin and immediate fate of the AVE

Following our diffusion analyses of the spatial differences in gene expression among AVE and emVE cells, we next compared AVE cells from E5.5 and E6.25 together, in order to understand the transcriptional changes within the AVE as it matures and acquires its cellular function in patterning the underlying epiblast. The ratio between spliced and unspliced mRNA can be used to estimate the transcriptional dynamics of differentiating cell populations. We therefore performed RNA velocity analysis (La Manno et al., 2018) with the scVelo implementation (Bergen et al., 2020, see Methods), to understand the relationships among AVE and emVE cells at E5.5 and E6.25. The velocities of the single cells were projected onto the first two diffusion components of a diffusion map of AVE and emVE cells, computed by integrating the two stages (see Methods and Supplementary Figure 5). The stream indicates a trajectory originating from the E5.5 emVE, transitioning *through* the E5.5 AVE, then the E6.25 AVE and ending in the E6.25 emVE (Figure 4A).

**Figure 4.**
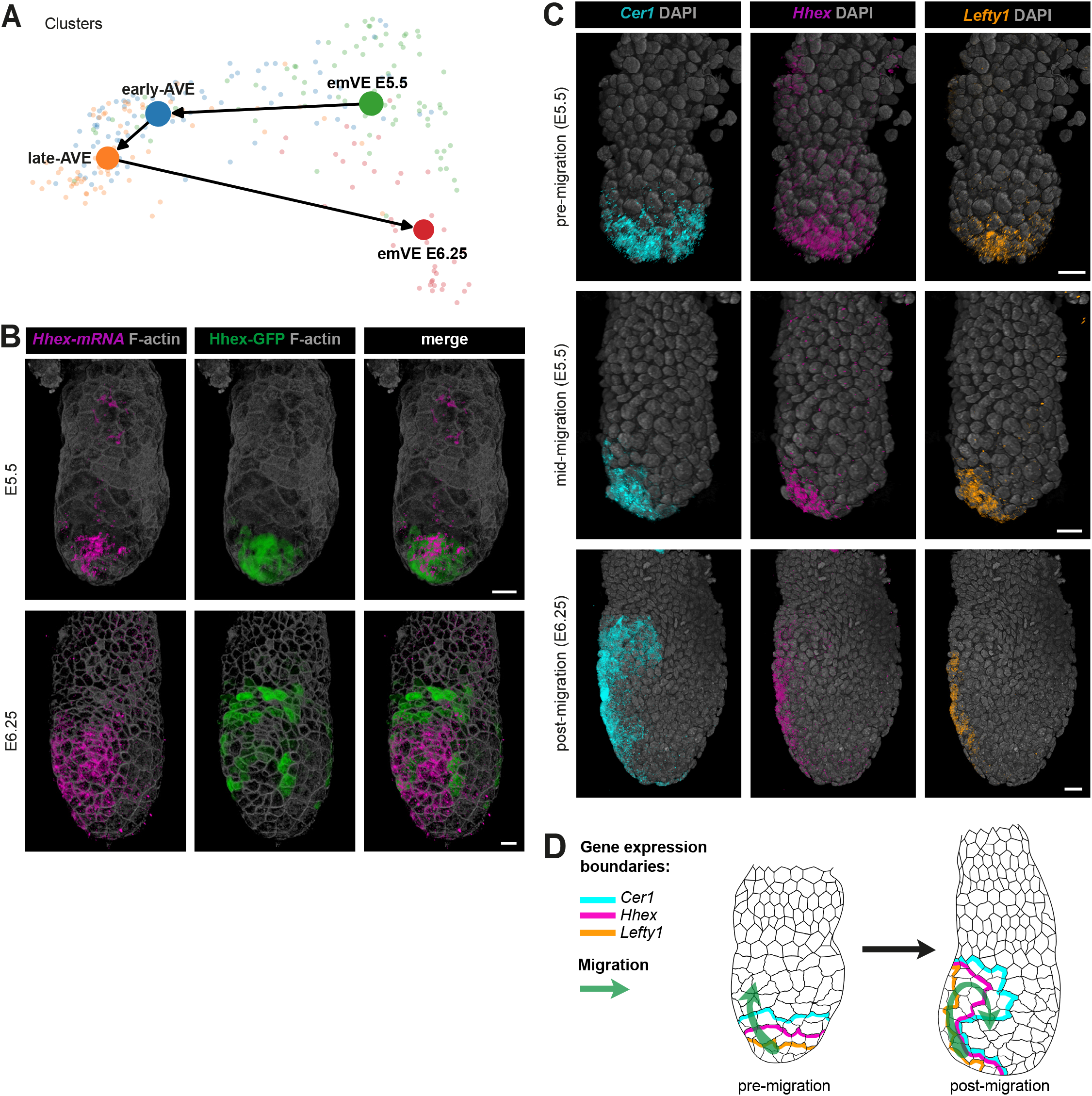
The origin and fate of the anterior visceral endoderm. (A) Partition-based graph abstraction (PAGA) graph computed from RNA velocities, and projected on the first two diffusion components of a diffusion map computed on the AVE and emVE clusters, combining the data from the stages E5.5 and E6.25. (B) Validation of the RNA velocity results by the direct comparison of short-term lineage labelled AVE cells (expressing the *Hhex-GFP* transgene reporter) and the contemporaneous expression of the endogenous *Hhex* gene (visualised by *in situ* HCR) in E5.5 and E6.25 embryos. (C) Volume renderings directly comparing the changes to the expression domains of the AVE-markers *Cer1, Hhex*, and *Lefty1* (visualised by *in situ* HCR) in embryos before, during and after AVE migration. (D) Schematic summarising the changes to the *Cer1*, *Hhex*, and *Lefty1* expression domains within the VE as a consequence of AVE migration. All scale bars represent 20 *μ*m and all embryos are orientated with the anterior on the left (←).

**Figure 5.**
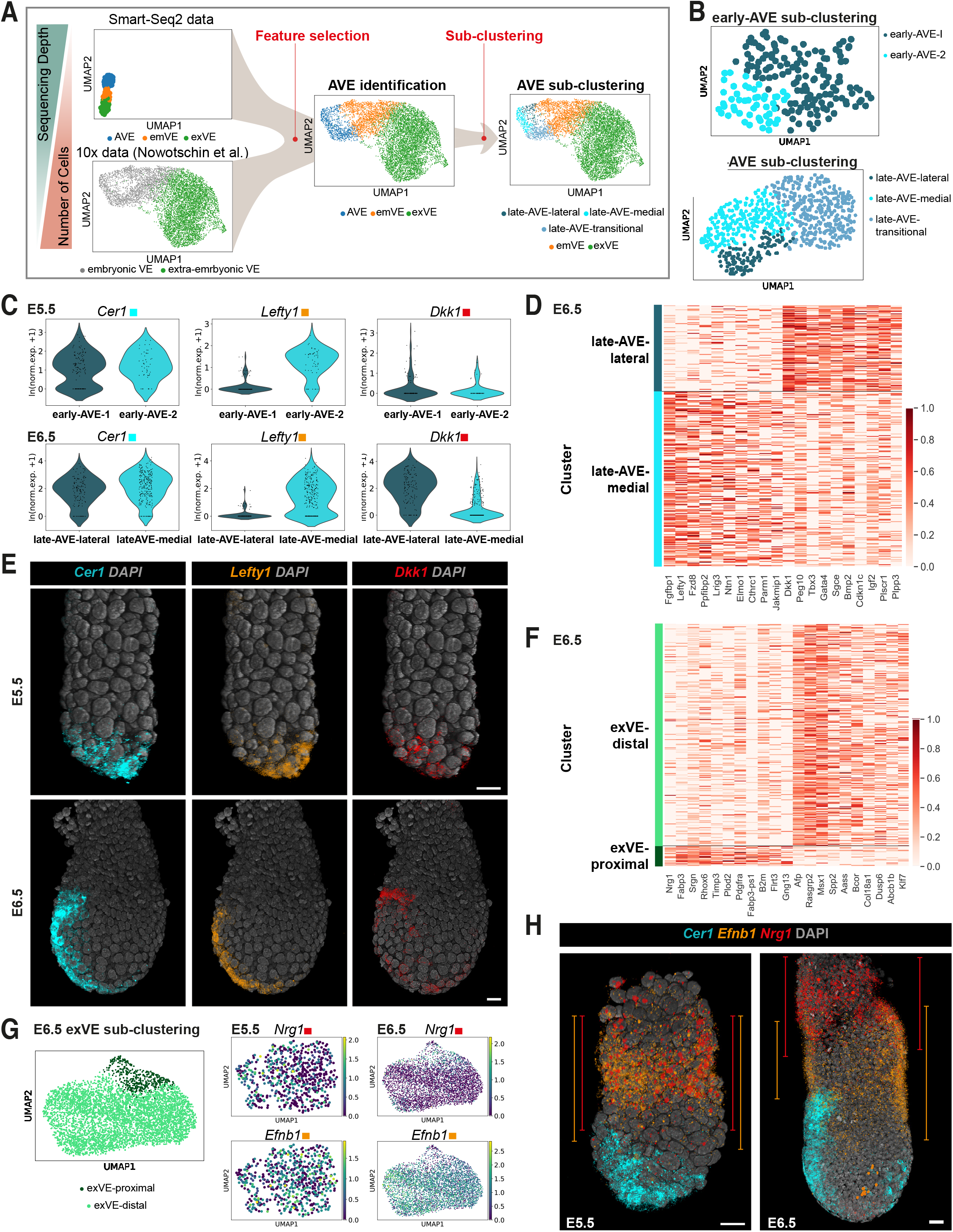
Relationship between the transcriptional heterogeneity in the AVE and exVE clusters and the spatial position of the cell sub-types in embryos. (A) Computational strategy used for the sub-clustering analysis of the scRNA-seq data from (Nowotschin et al., 2019), sequenced using the 10x protocol. Given the higher sequencing depth of our Smart-seq2 data, and with the goal of exploiting the larger number of VE cells present in the 10x dataset, we used the gene signatures obtained from the former for the identification of the AVE and emVE clusters in the 10x data. Then we selected the AVE cluster, and we performed a sub clustering analysis to dissect its heterogeneity (*see* the Methods section). (B) UMAP plots of the cells belonging to the AVE at E5.5 (top) and E6.5 (bottom), (data from Nowotschin et al., 2019). Cells are coloured according to the groups identified through a sub-clustering analysis (*see* the Methods section); *see* Figure S6 and the Methods section for AVE identification in these data. (C) Violin plots showing *Cer1*, *Lefty1* and *Dkk1* normalized log expression levels at E5.5 (top) and E6.5 (bottom). Cells are grouped according to the clusters shown in (B). (D) Heatmap showing the expression of the top 10 genes upregulated in the late-AVE-medial cluster and the top 10 genes upregulated in the late-AVE-lateral cluster, obtained through a differential expression analysis between the late-AVE-medial and the late-AVE-lateral clusters E6.5. The normalized log expression levels of each gene are standardised so that they vary within the interval [0,1]. (E) Volume renderings showing the expression domains of *Cer1, Lefty1*, and *Dkk1* (visualised by *in situ* HCR) at E5.5 and E6.5 used to distinguish between the sub-regions of the AVE. (F) Heatmap showing the expression of the top 10 genes upregulated in the exVE-proximal cluster and the top 10 genes upregulated in the exVE-distal cluster, obtained through a differential expression analysis between the exVE-proximal and the exVE-distal clusters at E6.5. The normalized log expression levels of each gene are standardised so that they vary within the interval [0,1]. Note that *Nrg1* (first gene shown on the left) was manually added to the list, since it was not in the top 10 genes upregulated in the exVE proximal cluster (it was the 13^th^ in the ranking by adjusted p-value). (G) UMAP plot of the cells belonging to the exVE cluster at E6.5 (left), coloured according to the groups identified through a sub-clustering analysis (*see* the Methods section); UMAP plots of the cells belonging to the exVE cluster at E5.5 (centre) and E6.5 (right), coloured according to the normalized log expression of the genes *Nrg1* and *Efnb1*. (H) Volume renderings showing the changes to the expression domains of *Efnb1* and *Nrg1* (visualised by *in situ* HCR) between E5.5 and E6.5 in comparison to that of *Cer1* (marking the position of the AVE). The orange and red lines represent, for *Efnb1* and *Nrg1* respectively, the proximal and distal extent of their expression domains. All scale bars represent 20 *μ*m and all embryos are orientated with the anterior on the left (←).

It was clear from both the diffusion map and the RNA velocity analysis that early- and late-AVE are not transcriptionally equivalent (Figure 4A and Supplementary Figure 5). This suggested the hypothesis that upon induction at E5.5, AVE cells (equivalent to the DVE) emerge as a transcriptionally distinct population from the emVE, becoming transcriptionally more divergent from it as they migrate anteriorly and ‘mature’, but then transition back toward an emVE state as they are displaced laterally by the following stream of cells that give rise to the AVE at E6.25. This highlights the transient nature of the AVE and further predicts that AVE cells, once displaced laterally, would lose markers characteristic of the AVE in reverting back to an emVE state.

We performed a short-term lineage labelling experiment to test this hypothesis, using the well-established *Hex-GFP* line (Nowotschin et al., 2013; Rakeman and Anderson, 2006; Rodriguez et al., 2001; Srinivas et al., 2004) that labels AVE cells with GFP from E5.5 to beyond E6.5. We took advantage of the perdurance of GFP protein to label cells that expressed *Hhex* in the past, combined with HCR to assay the current transcriptional status based on the presence or absence of the endogenous *Hhex* transcript (Figure 4B). At E5.5 *Hhex* mRNA and Hhex-GFP fluorescent signal both colocalised in migrating AVE cells at the distal end of the egg cylinder, verifying that Hhex-GFP is a good reporter of endogenous *Hhex* expression. However, by E6.25, in embryos in which migrating AVE cells had reached the boundary between the emVE and exVE, and had started to be displaced laterally, there was less congruence between the expression of endogenous *Hhex* mRNA and the Hhex-GFP reporter. The most proximal and lateral cells marked by Hhex-GFP expression no longer expressed endogenous *Hhex* transcript (Figure 4B) indicating that cells that had expressed the archetypal AVE marker *Hhex* in the past now no longer do so as they move laterally. This is consistent with the RNA velocity analysis that AVE cells at E6.25 revert to an emVE state as they get displaced laterally.

To further test this hypothesis, we used HCR to look at the expression of multiple AVE markers (*Cer1* and *Lefty1*) characteristic of the AVE signature, along with *Hhex*, in the same embryos at different stages of migration. Such multiplexing is important in mitigating against inconsistencies that otherwise arise from variations across embryos if using different embryos for different markers at a stage when transcriptional, positional, and cellular changes occur rapidly, over a very short time-period. We found that before migration is initiated, the expression of *Cer1* extended proximally over a larger region of the AVE in comparison to *Lefty1*, which was restricted to the distal-most cells (N=13; Figure 4C). At mid-migration stages, we detected a discernible asymmetry in the expression domains of all three transcripts, with the *Cer1* expression domain always being the widest and most proximal in extent, followed by that of *Hhex*, and then *Lefty1* (N=18). At post-migration stages, the domain of expression of *Lefty1* and *Hhex* were never seen to extend laterally, while *Cer1* was detected at only relatively low levels in the more lateral cells (N=8), consistent with a successive loss of AVE identity, and a gradual shift back towards an emVE state as AVE cells get displaced laterally (Figure 4D).

### The AVE is a spatially and temporally heterogeneous cell population

Our scRNA-seq analysis showed that the expression of *Cer1*, *Lefty1* and *Hhex* is heterogeneous within the AVE (Figure 1C), while the HCR experiments revealed how the observed heterogeneity was linked to the spatial position of the cells within the migrating AVE (Figure 4C and D). This suggested that even within the relatively small group of AVE cells in an embryo, there are transcriptionally distinct sub-populations of cells that may also be spatially distinct.

To uncover these AVE sub-populations, we took advantage of the large numbers of cells in a recently published scRNA-seq dataset obtained using the 10x protocol, which includes 1,067 cells from the VE at E5.5 and 7,182 VE cells at E6.5, annotated as embryonic or extra-embryonic visceral endoderm (Nowotschin et al., 2019). We leveraged the higher sequencing depth of our Smart-seq2 dataset, which allowed us to accurately define the transcriptional signatures for different AVE cell types, with the large number of cells in the 10x dataset that, despite the shallower sequencing depth, allowed the identification of cell sub-clusters with greater confidence. We first used the information from our Smart-seq2 dataset to identify the AVE cells from amongst the embryonic visceral endoderm cluster of the 10x dataset at E5.5 and E6.5 (see Methods, Supplementary Figure 6A and B). We then sub-clustered the AVE cells at the two stages (see Methods and Figure 5A).

Three AVE sub-clusters emerged at E6.5 from the 10X dataset (Figure 5B ‘late-AVE sub-clustering’ and Supplementary Table 9). One of these clusters, despite having low levels of several canonical AVE-markers including *Cer1, Hhex*, and *Lefty1* (Supplementary Figure 6C), in comparison to emVE cells, showed relatively higher levels of the ‘high-in-AVE’ genes we identified previously (Figure 1D, Supplementary Figure 6D and Supplementary Table 10). This cluster therefore likely represented a transitional state (late-AVE-transitional) occupied by cells that originated as the early-AVE at E5.5 and are in the process of downregulating the AVE-transcriptional program to acquire an emVE-like transcriptional state. This was also consistent with our HCR results for expression of *Cer1, Hhex*, and *Lefty1* and our AVE fate analysis (Figure 4B and C). The two other sub-clusters both had high *Cer1*-expression and were distinguished by the expression of *Lefty1* in one and *Dkk1* in the other (Figure 5C) along with other differentially expressed genes (Figure 5D and Supplementary Table 11). Using multiplexed-HCR for *Cer1*, *Lefty1* and *Dkk1* to provide spatial landmarks, we mapped these two sub-clusters of the late-AVE onto the medial and lateral regions of the E6.5 embryo respectively (Figure 5E, ‘E6.5’).

While we could not find sub-clusters in AVE at E5.5 with an unsupervised approach (see Methods), to determine whether the AVE sub-clusters identified at E6.5 are already present in the AVE at E5.5, we next performed a sub-clustering of E5.5 AVE cells from the 10X dataset, employing the top 20 genes differentially expressed between the AVE-medial and -lateral sub-clusters at E6.5 (Figure 5B, ‘early-AVE sub-clustering’). The AVE sub-clusters found through this analysis at E5.5 had comparable levels of *Cer1* but different levels of *Lefty1* (Figure 5C). HCR shows that these two sub-populations are spatially segregated at E5.5 with the *Cer1*+, *Lefty1*+ population (early-AVE-1) located at the very distal tip of the embryo and *Cer1*+, *Lefty1*-cells (early-AVE-2) more proximally (Figure 5E, ‘E5.5’). However, the transcriptional differences between these two sub-clusters are very limited (with *Lefty1* and *Fgfbp1*, being the only differentially expressed genes), suggesting that heterogeneity is a feature which the AVE cells acquire only as they undergo their characteristic migration to the prospective anterior and is related to the spatial positions the cells occupy within the embryos.

Finally, we used this approach to see if any additional heterogeneities were present amongst the exVE cells, that ultimately contribute to the Yolk Sac endoderm. At E6.5, two transcriptionally distinct sub-clusters emerged, one marked by *Nrg1* and the other by *Efnb1* (Figure 5F and G; Supplementary Table 12), both genes we previously identified as markers of the non-migratory emVE (Figure 1F). At E5.5 however, these two markers did not distinguish transcriptionally distinct sub-populations within the exVE (Figure 5G). To verify whether these transcriptional clusters showed spatial segregation in the embryo, we detected *Nrg1* and *Efnb1* at E6.5 using HCR.

This revealed that the two populations were spatially segregated, with *Nrg1*+, *Efnb1*-cells located more proximally and enriched in the ectoplacental cone in comparison to *Nrg1*-*Efnb1*+ cells that were present in the distal exVE and the proximal emVE (Figure 5H). Consistent with the computational analysis, at E5.5, these markers showed significant overlap, indicating that heterogeneity within the exVE only emerges between E5.5 and E6.5.

### Ephrin- and Semaphorin-signalling pathways are important for the migration of the AVE

Migration of the AVE is central to its function but the processes controlling the initiation and directionality of migration are still unknown. We reasoned that signalling interactions between the AVE and surrounding tissues might provide cues to initiate and drive AVE migration, and therefore sought to identify possible signalling interactions from our scRNA-seq dataset. To do this, we started with a list of 2,548 ligand-receptor pairs (LRPs) (Caruso et al., 2020) that we filtered down to those most likely to mediate cell-cell communication in our dataset (see Methods). Among the resulting LRPs, we then considered those containing genes from the ‘high-in-AVE’ groups we identified from the diffusion pseudotime analysis (Figure 1D). Finally, we employed COMUNET (Solovey and Scialdone, 2020) for the visualisation and exploration of possible ligand-receptor interactions between the various cell types of the embryo at E5.5 and E6.25.

We noticed that many of the putative interactions identified involved the bidirectional Eph/Ephrin-signalling pathway (Figure 6A). Moreover, even just a subset of the genes belonging to this pathway was sufficient for separating all the five cell clusters previously identified at E5.5 and E6.25 using the entire transcriptome (Figure 6B, Supplementary Figure 7A–C and Methods), suggesting that Eph/Ephrin-signalling might play cell-type-specific roles at these stages. *Efna5* and *Ephb3* in particular showed relative higher expression in the AVE compared to emVE or epiblast (Supplementary Figures 8A and 9A, C). We therefore plotted potential communication patterns for *Efna5* and *Ephb3* at E5.5, that revealed several plausible signalling interactions between the AVE and adjacent epiblast or emVE cells (Supplementary Figure 8B), raising the possibility that these short-range intercellular interactions might regulate AVE migration in some way.

**Figure 6.**
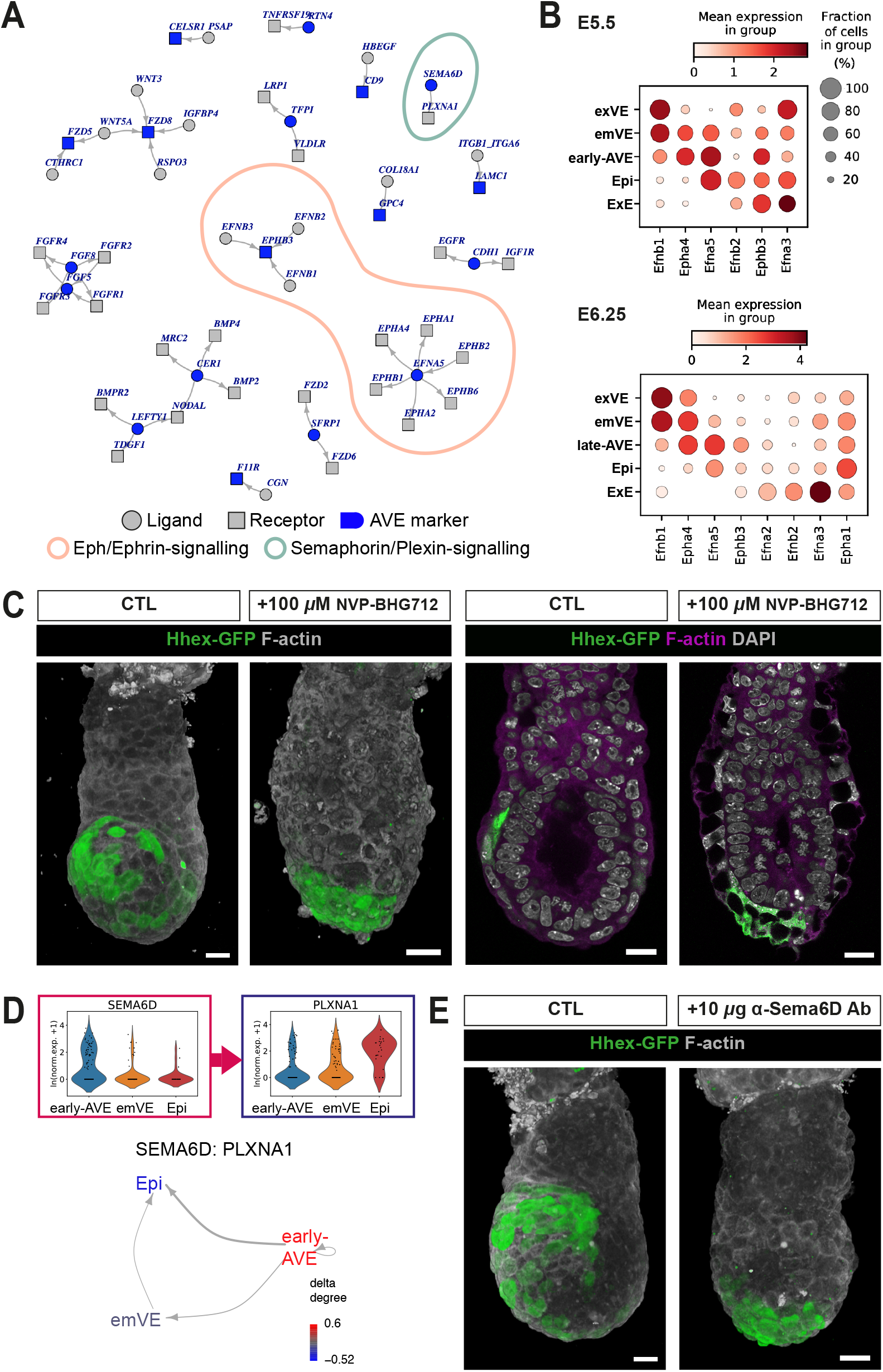
Ephrin- and Semaphorin-signalling are important for AVE migration. (A) Graph showing ligands and receptors belonging to Ligand–Receptor Pairs (LRPs) containing at least one ‘high-in-AVE’ gene, identified from the diffusion pseudotime analysis, at E5.5 and E6.25. Circles and squares indicate ligand and receptors, respectively, and AVE-markers (from the ‘high-in-AVE’ group) are coloured in blue. The direction of the arrows indicates whether the gene is acting as a ligand or a receptor in a given LRP. The red shaded area indicates LRPs belonging to the Eph/Ephrin-signalling pathway, the green shaded area indicates the LRP formed by SEMA6D and PLXNA1, belonging to the Semaphorin/plexin signalling pathway. (B) Dot plots of genes belonging to the Eph/Ephrin signalling pathway at E5.5 (top) and E6.25 (bottom). The genes were selected as those showing a significant contribution to the first three principal components in a Principal Components Analysis performed on genes belonging to the Eph/Ephrin signalling pathway (see Methods and Figure S7). (C) Visualisation of AVE migration in embryos expressing the *Hhex-GFP* transgene reporter, cultured with (+100 *μ*M NVP-BHG712) and without (CTL) the presence of the Eph/Ephrin-signalling inhibitor. The two images on the left are volume renderings showing the overall position of the AVE at the end of the culture period; the two images on the right are optical sections through the embryos showing the effects of the inhibitor on other cell types of the embryo including the emVE, exVE, and epiblast. (D) Violin plots showing the normalized log expression levels of *Sema6d* and *Plxna1* in the AVE, emVE and Epi clusters, at E5.5 (top), and the intercellular communication pattern associated to the LRP *Sema6d-Plxna1*. (E) Volume renderings visualising AVE migration in embryos expressing the *Hhex-GFP* transgene reporter, cultured in the presence of antibodies against Sema6D or PlexinA1 in comparison to a control (CTL) embryo. All scale bars represent 20 *μ*m and all embryos are orientated with the anterior on the left (←).

Eph receptors and their ligands, the Ephrins, have been implicated in various developmental processes such as tissue-boundary formation, cell adhesion, migration and axon guidance (Park and Lee, 2015) but have not been implicated to play a role in AVE migration. To test if they were indeed involved in AVE migration as suggested by our intercellular communication analysis, we used pharmacological blockade of Eph/Ephrin-signalling in E5.5 embryos expressing the *Hex-GFP* reporter to monitor AVE migration. Control cultured embryos showed migration of AVE cells from the distal tip to the emVE–exVE margin and then laterally across the epiblast (N=25/29). However, embryos cultured in the presence of the broad-spectrum Eph/Ephrin-signalling inhibitor NVP-BHG712 (Martiny-Baron et al., 2010; Tröster et al., 2018) showed a failure of AVE cells to properly migrate from the distal tip (N=29/32) (Figure 6C). They also had more mitotic cells within the epiblast, and abnormal nuclear morphology in the VE, indicating that, consistent with our analyses, Eph/Ephrin-signalling in addition to its effect on cell migration, plays a variety of important cell-type specific roles in the E5.5 embryo.

Another ligand-receptor pair identified by our intercellular communication analysis included the bi-directionally signalling trans-membrane proteins SEMA6D and PLXNA1 (Figure 6A, D). We selected these for further study on the basis of *Sema6d’s* association with the GO-term ‘cell migration’ and lack of a previously reported role in the pre-gastrulation embryo. *Sema6d* was expressed at higher levels in the AVE, while its known receptor, *PlxnA1*, was expressed in the surrounding emVE cells as well as the epiblast (Supplementary Figure 9B, D). To test if SEMA6D also has a role in AVE migration, we cultured E5.5 embryos expressing the *Hex-GFP* transgene, in the presence of an inhibitory antibody against SEMA6D (Figure 6E). Control embryos showed AVE migration as expected (N=18/20), but embryos cultured with the inhibitory antibody showed an arrest in migration (N=21/26), consistent with a novel role for Semaphorin-signalling in AVE migration.

## DISCUSSION

The collective directional migration of the AVE is one of the first such migration events to occur in the developing mouse embryo (Srinivas et al., 2004; Thomas and Beddington, 1996). It plays a pivotal role in specifying the first definitive axis of the body, the A–P axis, upon which all future development is predicated. In this study, by coupling complementary experimental techniques, such as scRNA-seq, high-resolution imaging, multiplexed-HCR and short-term lineage labelling, we reveal important new insights into the dynamic nature of this small population of cells over a short period of developmental time.

Our single-cell analysis reveals considerable transcriptional heterogeneity within the VE of the pre-gastrulation embryo. We find that by E6.25, the exVE consists of two transcriptionally distinct sub-groups of cells, which are also spatially segregated along the proximal– distal axis. At this stage, the AVE can be sub-divided into three transcriptionally distinct cell types (medial, lateral, and transitional), that again show spatial segregation. This reveals that the VE is spatially patterned to a remarkable degree already at this early a stage of development compared to the rest of the embryo, despite representing a relatively small field of cells.

Using GFP as a short-term lineage label and mRNA expression as a readout of the current transcriptional state of cells, we validated a prediction made by our RNA-velocity analysis, that the E5.5 AVE cells—that originate at the distal tip but occupy a more lateral position upon completing migration—downregulate their AVE-specific transcriptional programme and revert to a more general emVE transcriptional state. This gradual attenuation of ‘anteriorising’ gene expression as AVE cells migrate laterally is necessary to restrict the inhibition of WNT and NODAL signalling to the anterior of the egg cylinder while allowing the formation of the primitive streak in the posterior half. Such tight coupling of transcriptional state and position is presumably required because of the constraints of the relatively small egg cylinder, within which patterning has to be achieved in a precise and reproducible manner. This also indicates that the AVE at E5.5 and E6.5 albeit being broadly the same cell-type, capture different transcriptional states of an everchanging population and exhibit different degrees of heterogeneity which can be interpreted as different levels of ‘maturity’. The transcriptional state observed among the E6.5 AVE (the most transcriptionally distinct and mature), is thus a transient one, acquired by migrating VE cells and is directly related to the position the cells occupy on the embryo at any given time.

The migration of the AVE cells and the change in their transcriptional state both ultimately lead to gross asymmetries in gene expression within the VE along the A– P axis. In contrast to previous reports (Rivera-Pérez and Magnuson, 2005), we show that asymmetries in gene expression are manifest first in the AVE and only subsequently in the posterior regions of the embryo. This difference in results is likely due to the great variation in morphology across embryos at this stage (Lawson and Wilson, 2016) with changes occurring over a short period of time, that confound attempts to stage-match embryos for the purpose of comparing gene expression dynamics. Comparing multiple transcripts in the same embryo via multiplexed-HCR overcomes such problems. These findings again suggest that transcriptional changes within the AVE are coordinated closely with migration over relatively short time-scales, so that these cells can correctly and precisely pattern the underlying epiblast. Our HCR experiments indicate that at this stage, the epiblast is only starting to show evidence of patterning, in the form of asymmetric expression of genes such as *Nodal* and *T*, reinforcing the notion that *in utero*, axial asymmetry is established first in the VE and only subsequently in the epiblast.

Comparison of phosphoproteomic with transcriptomic data identified AVE-enriched proteins such as DBN1, MARCKS and MARCKSL1, that are known to control the morphology and motility of a wide range of cell-types (El Amri et al., 2018; Shirao and Sekino, 2017), consistent with a potential role in AVE migration. In particular, DBN1 plays an important role in the regulation of actin filament reorganization (Grintsevich, 2021; Tanabe et al., 2014). Actomyosin dynamics is also known to play a role in AVE migration with sub-cortical F-actin being enriched in a ring delineating the apical junctions of emVE cells (Trichas et al., 2011) and regulators of F-actin branching being essential for normal AVE migration. Other proteins such as PML and KMT2A, which are known to be involved in the regulation of cell migration (Amodeo et al., 2017; Zhang et al., 2017), were shown by the transcriptomic data to be expressed throughout the VE, but were found to be differentially phosphorylated specifically in the embryonic half, which might point to them playing a role in AVE migration, following post-translational modification. Overall, leveraging phosphoproteomic data against transcriptomic data in this way can help narrow down interesting candidates to pursue for future functional studies.

Distinct from the cell-autonomous mechanistic bases for AVE motility suggested by the above approach, our communication analysis and inhibitor experiments identified novel intercellular signalling pathways that might be involved in the unidirectional migration of the AVE. Ephrins, Semaphorins and their respective cognate receptors are mainly membrane-tethered molecules, that were first identified as axon guidance cues, but have since been shown to regulate migration of other cell types by direct intercellular signalling (Alto and Terman, 2017; Chen et al., 2021; Kania and Klein, 2016; Toyofuku et al., 2004). Given the large number of Ephrin and Semaphorin family members expressed in the early embryo at this stage, a certain degree of functional redundancy is possible. Furthermore, the bidirectional nature of their signalling, makes it likely their precise roles in modulating AVE migration might be complex. However, recently it has been shown that in the follicular epithelium of *Drosophila* egg chambers, transmembrane Semaphorins can be planar polarised and enriched at the leading edge of the basal surface of cells, from where they coordinate unidirectional collective migration through communication with the cell ahead expressing the corresponding Plexin receptor (Stedden et al., 2019). We could hypothesise a similar mechanism, through which SEMA6D in the migrating AVE cells acts as a repulsive cue for the surrounding emVE cells expressing the PLXNA1 receptor, while playing AVE-autonomous roles to ensure collective unidirectional motility. Similarly, Eph– Ephrin interactions can regulate the directional nature of cell movements through coordinating contact inhibition (Astin et al., 2010; Batson et al., 2013). They are also necessary for boundary formation between cells sharing a common plane or border (Cayuso et al., 2015). The differentially-expressed *Eph* and *Ephrins* we have identified within the cell types of the early embryo, in addition to facilitating AVE migration could be postulated to form the molecular basis for the emVE–exVE boundary that marks the proximal extent of AVE migration.

Taken together, these findings highlight the power of utilising transcriptomic data at a cellular and temporal resolution alongside careful experimental validation to further our understanding of highly dynamic cell-populations such as the AVE, which migrates and changes its transcriptional identity in a coordinated manner. Our data shows that this dynamic nature of the AVE, overtly observed as its migratory behaviour and transient function, emerges from an elaborate series of continually changing molecular heterogeneities underpinning it.

## METHODS

### Mouse Strains, Husbandry and Embryo Collection

All animal experimentation procedures were performed in full accordance with the UK Animals (Scientific Procedures) Act 1986, approved by Oxford University’s Biological Services Ethical Review Process and were performed under UK Home Office project licenses PPL 30/3420 and PCB8EF1B4. Mice were maintained on a 12 h light, 12 h dark cycle and the noon on the day of finding a vaginal plug was designated 0.5 dpc (E0.5). Embryos were collected at the appropriate stages between E5.25-E6.5, from C57BL/6J (in house) or CD1 (Charles River, England) females crossed to C57BL/6J or homozygous *Hex-GFP* transgenic studs (Rodriguez et al., 2001). The dissections were done according to a standard post-implantation dissection protocol as previously described Srinivas et al. (2010). All dissecting instruments were thoroughly cleaned with RNase*Zap*™ (Invitrogen, AM9760) and 70% ethanol, and embryos being collected for HCR were kept in ice-cold M2 medium (Sigma-Aldrich, M7167) throughout.

### Single-cell isolation, cDNA library preparation and sequencing

Embryos were collected at E5.5 (N=40) and E6.25 (N=11) from C57BL/6J females crossed to C57BL/6J studs. Fluorescent membrane labelling of the VE was achieved using the CellVue Claret Far Red Fluorescent Cell Linker Kit (Sigma-Aldrich, MINICLARET-1KT). Briefly, embryos were incubated in 0.1% (v/v) Claret Far Red dye in Diluent-C for 5 minutes at room temperature (RT); the reaction was stopped with an equal volume of 1% BSA, and then rinsed with M2 medium. Up to 4 stained embryos were placed in 100 *μ*l of TrypLE™dissociation reagent (Invitrogen, 12563011) for 3.5 minutes and 4.5 minutes for E5.5 and E6.25 embryos respectively, at 37°C. The embryos were then mouth-pipetted up and down, for gentle mechanical dissociation using a glass capillary 10-15% larger than the size of the embryos. The dissociated cells were pooled and transferred to a 1.5 mL microcentrifuge tube and the TrypLE was neutralised with an equal volume of heat-inactivated FBS (Thermo Fisher, 10500) followed by centrifugation at 1000x *g* for 3 minutes at 4°C. The cells were resuspended in 100 *μ*l of ice-cold HBSS (Sigma-Aldrich, 55037C) with 1% FBS. DAPI (0.1 *μ*g/mL; Vector Labs, H-1200) was added as a live-dead indicator. Claret-labelled, live, VE cells were collected using SH800 Cell Sorter (Sony Biotechnology) directly into plates containing lysis buffer at 4°C.

Total mRNA from the cells were extracted and amplified using the SMARTSeq2 protocol (Picelli et al., 2014) with the additional inclusion of ERCC spike-in control at 1/10^7^ concentration. Multiplexed sequencing libraries were generated from cDNA using the Illumina Nextera XT protocol. 125 bp paired-end sequencing was performed on an Illumina HiSeq 2500 instrument (V4 Chemistry).

### Transcript quantification, quality control and normalization

We performed transcript quantification in the scRNA-seq datasets at stages E5.5 and E6.25 employing Salmon (v0.13.1) (Patro et al., 2017) in the quasi-mapping-based mode. First, we created a transcriptome index from the mouse reference (version GRCm38.p6) and ERCC spike-in sequences. Then, we used the “quant” function to quantify the transcripts, correcting for the sequence-specific biases (“--seqBias” flag) and the fragment-level GC biases (“--gcBias” flag). Finally, we aggregated the transcript level abundances to gene level counts. The obtained raw count matrices include 384 samples at each stage (E5.5 and E6.25).

Afterwards, we performed a quality control to eliminate low quality cells from downstream analyses. We selected good quality cells according to the following criteria (same for E5.5 and E6.25): Number of genes with more than 10 reads per million (rpm) larger than 3,000;

– Log10 of the total number of reads larger than 4;
– Fraction of mapped reads larger than 0.5;
– Fraction of reads mapped to mitochondrial genes smaller than 0.1;
– Fraction of reads mapped to ERCC spike-ins smaller than 0.3.

With these criteria, we obtained 255 good quality cells at E5.5 and 238 cells at E6.25.

We normalized the raw count matrices separately at E5.5 and E6.25 using the R package “scran” (v1.10.2) (Lun et al., 2016) with default parameters, and we log-transformed the data (adding a pseudocount of 1 in order to avoid infinities) using the natural logarithm (function “scanpy.pp.log1p” in Scanpy v1.4 (Wolf et al., 2019)).

### Cell clustering and assignment of cluster identities

We performed hierarchical clustering of the cells from each stage separately, using an information theoretic criterion to guide the choice of the number of clusters, as described below. First, we computed the highly variable genes (HVGs) employing the Scanpy function “scanpy.pp.highly_variable_genes”, with default parameters except for “max_mean” (set to 10), and retained the top 3,000 genes at both E5.5 and E6.25 stages.

Then, we computed the distance matrix between cells as 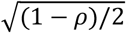 where *ρ* is the Spearman’s correlation coefficient between cells. Hierarchical clustering was carried out on this distance matrix (function “hclust” in R, with average agglomeration method) and we cut the dendrogram with the dynamic hybrid cut method (Langfelder et al., 2007) (“cutreeDynamic” function in the R package “dynamicTreeCut” v1.63.1, with the hybrid method and a minimum cluster size of 10 cells). This method depends on the parameter “deepsplit”, which ranges between 0 and 4 and determines the number of clusters.

To estimate the number of clusters, for each value of “deepsplit”, we computed the average Variation of Information (Meilă, 2007) between the clustering obtained using the top 3,000 HVGs and 50 subsamples in which only half of the genes is randomly kept. Similar to the elbow method, we chose the largest “deepsplit” value before the average Variation of Information has a ‘kick’ towards high values (see for instance Supplementary Figure 1B). The Variation of Information is computed using the function “vi.dist” in R package “mcclust” v1.0 (Fritsch and Ickstadt, 2009). This procedure gave 4 clusters at E5.5 and 3 clusters at E6.25.

At E5.5, two clusters corresponded to Visceral Endoderm (VE) cells, which, based on their marker genes, could be identified as the VE portions covering the extraembryonic ectoderm (exVE) and the epiblast. We further divided the latter into two clusters by recomputing the HVGs and by employing the “Partitioning Around Medoids” (pam) function (R package “cluster” v2.1.0). The function was run on the Spearman’s correlation distance matrix, computed as described above. This allowed us to distinguish an AVE cluster from the rest of the VE covering the epiblast (emVE). At E6.25, while one cluster shows a clear AVE signature, the other two include multiple cell populations, as can be seen from the expression of marker genes. In particular, the first includes exVE and emVE together, which we separated by recomputing the HVGs and by applying the “pam” function as described above. The second cluster mostly includes epiblast cells, while a few cells express extraembryonic ectoderm (ExE) markers. We identified the ExE cells using an outlier detection algorithm based on the distance from the k-nearest neighbours for each cell (python package “PyOD: https://pyod.readthedocs.io/en/latest/index.html) (Zhao et al., 2019), function “KNN” in “pyod.models.knn”).

Overall, we identified 5 clusters each in E5.5 and E6.25 embryos: one corresponding to epiblast cells (Epi), another including cells from the extra-embryonic ectoderm (ExE) and three clusters of visceral endoderm cells (emVE, exVE and AVE). Three cells at each stage were unassigned in the clustering analysis; we eliminated them for downstream analyses, ending up with 252 cells at E5.5 and 235 at E6.25.

We computed markers for the clusters relying on the Scanpy function “scanpy.tl.rank_genes_groups”. For each pair of clusters, we tested the differential expression of genes using the Wilcoxon test, with Benjamini-Hochberg correction for multiple testing. We selected genes with log2 fold change larger than 1 and adjusted p-value smaller than 0.1. For each cluster, we ranked the genes based on their average -log10 of the adjusted p-values across all pairwise comparisons. The heatmaps (Supplementary Figure 1) were generated considering the top five markers per cluster. Note that some genes can be markers of more than one cluster. The top 50 markers per cluster at the two stages are listed in Supplementary Tables 1-2. The UMAPs at E5.5 and E6.25 are generated using the Python package “umap” v1.3.9 (Becht et al., 2019), with 30 nearest neighbours using the same distance matrix that was used for clustering.

For the computation of the relative distances between VE cluster centroids (Supplementary Figure 1B), we identified the centroids using the “NearestCentroid” function in the Python package “scikits-learn” v0.21.3. Then, we computed the Spearman’s correlation distance between them, as described above. The PAGA graphs for the VE clusters at the two stages (Supplementary Figure 1B), were computed with the Scanpy function “scanpy.tl.paga” (Wolf et al., 2019).

### Diffusion pseudotime analysis of AVE and emVE cells

We selected only AVE and emVE cells at each stage and we identified the top 3,000 HVGs as described above. Using these genes, we performed a Principal Component Analysis (PCA) (Scanpy function “scanpy.tl.pca”, with “arpack” as SVD solver) and we built a k-nearest neighbour (knn) graph of the cells (Scanpy function “scanpy.pp.neighbors” with k=15) based on the Spearman’s correlation distance calculated on the first 10 Principal Components.

Starting from this knn graph, we computed a diffusion map (Scanpy function “scanpy.tl.diffmap”) and a pseudotime coordinate (Scanpy function “scanpy.tl.dpt”), choosing as the root cell the one with minimum and maximum value of the second Diffusion Component (DC) at E5.5 and E6.25, respectively.

To find genes differentially expressed in pseudotime, first we filtered out genes detected in fewer than 10 cells. Then, we employed a Generalized Additive Model (R function “gam” from “GAM” package v1.16.1) to fit the expression in pseudotime of each gene and we computed a p-value using the ANOVA test for parametric effects provided by the “gam” function. After FDR correction, we obtained 952 differentially expressed genes at E5.5 and 915 at E6.25 (FDR < 0.01).

We classified the differentially expressed genes based on their trend in pseudotime. To do so, we clustered genes with the same approach used for cell clustering, but this time we fixed a minimum cluster size of 50 and the random samples for the computation of the average Variation of Information were obtained by randomly sampling 70% of cells from the dataset 50 times. We identified two groups of differentially expressed genes at each stage (with deepsplit = 2 and 1 at E5.5 and E6.25, respectively), one of genes with decreasing expression in pseudotime (“high-in-AVE genes”) and the other with increasing expression in pseudotime (“low-in-AVE genes”).

The lists of the differentially expressed genes with gene group assignment at each stage are reported in Supplementary Tables 3-4.

### Proteomics and phosphoproteomics analysis

A total of 104 embryos were dissected at E6.5 as previously described (Srinivas, 2010). Using fine tungsten needles, the embryos were carefully bisected along the epiblast–ExE boundary and the embryonic and abembryonic halves generated were pooled separately. The embryonic half (EPI half) included the epiblast and the visceral endoderm surrounding it; the abembryonic half (ExE half) had the extra-embryonic ectoderm and the associated visceral endoderm cells (see Supplementary Figure 1A). Sample preparation for proteomics was carried out as previously described in Casado et al. (2013). Cells from both pools were harvested, lysed and treated with phosphatase inhibitors. Each pool was further divided into four aliquots that were processed individually as technical replicates. Further treatment on resultant peptide solutions included enrichment of phosphopeptides via Immobilized Metal Ion Affinity Chromatography (IMAC; Alcolea et al., 2009). We analysed the phosphoproteomes using liquid choromatography-tandem MS (LC-MS/MS) as previously described (Casado and Cutillas, 2011). Briefly, phosphopeptide pellets were resuspended in 20 *μ*l of 0.1% TFA, and 4 *μ*l was loaded into an LC-MS/MS system, which consist of a nanoflow ultrahigh pressure liquid chromatography (UPLC, nanoAccuity Waters) coupled online to an Orbitrap XL mass spectrometer (ThermoFisher Scientific). Each sample was run three times and the data for each sample was averaged over the three runs, to generate chromatogram data.

In the proteomics and phosphoproteomics experiments, 1,690 and 1,770 peptides were quantified, respectively (see Supplementary Tables 13-14 for the raw counts). We aggregated the peptide data to the protein level by summing the counts of peptides corresponding to the same protein. The data was normalized using the “normalyzer” function from the R package “NormalyzerDE” v1.0.0 (Willforss et al., 2019), which compares several quality metrics for different normalization methods. Following the analysis presented in Chawade et al. (2014), we chose the Loess method. For each sample, we averaged the expression levels of the proteins over the three runs.

We performed a differential expression test between the ExE and Epi halves for all the proteins in each dataset, using the function “normalyzerDE” from the same package mentioned above. For each protein, the function returns the log2 fold change between the ExE and Epi halves and the adjusted p-value.

We compared the proteomics and phosphoproteomics results by plotting the log2 fold changes in a scatter plot (Supplementary Figure 2D), for the proteins quantified in both datasets. Proteins are marked as differentially expressed if the log2 fold change is larger than 1 (in absolute value) and the adjusted p-value is smaller than 0.1. Proteins are labelled as “high-in-AVE” if the corresponding gene was found in the “high-in-AVE genes” group from the diffusion pseudotime analysis of the scRNA-seq data from AVE and emVE cells at E6.25 (see above). In the scatter plot, we also distinguish proteins with only one peptide in the dataset from those with multiple peptides.

To compare the proteomics and phosphoproteomics data with the scRNA-seq data, the cells at E6.25 were split into two groups, based on whether they belong to the “Epi half” (Epi, emVE and AVE cells) or the “ExE half” (ExE and exVE). After removing the genes detected in fewer than 10 cells, we identified the differentially expressed genes between these two groups of cells with the R package “DESeq2” (Love et al., 2014).

We represented the results in two scatter plots, showing the estimated log2 fold change in the scRNA-seq data on the *x*-axis and the log2 fold change in the proteomics (Supplementary Figure 2E) or phosphoproteomics data (Supplementary Figure 2F) on the *y*-axis. Genes/proteins are marked as differentially expressed if the log2 fold change is larger than 1 (in absolute value) and the adjusted p-value is smaller than 0.1. As before, genes/proteins are labelled as “high-in-AVE” if the corresponding gene was found in the ‘high-in-AVE’ genes group from the diffusion pseudotime analysis of the scRNA-seq data from AVE and emVE cells at E6.25 (see above).

### RNA velocity of AVE and emVE cells

Firstly, we selected AVE and emVE cells from the E5.5 and E6.25 datasets. Then, we integrated the data from the two stages using the “mnn_correct” function in the “mnnpy” Python package (the method was introduced in (Haghverdi et al., 2018); the Python implementation is available at https://github.com/chriscainx/mnnpy), using the intersection of the top 3,000 HVGs from the two stages. We scaled the data to zero mean and unit variance (Scanpy function “scanpy.pp.scale”, with max_value=10), and we computed a diffusion map with the same procedure as above (with 20 principal components and k=15).

To perform the analysis of RNA velocity, we generated the raw count matrices of spliced and unspliced counts at E5.5 and E6.25, using “STAR” v2.7.0f_0328 and “velocyto” v0.17.17, in “run-smartseq2” mode. We merged the matrices and performed the downstream analysis using the Python package “scvelo” v0.2.2 (Bergen et al., 2020) as described below. We filtered out genes expressed in fewer than 10 cells, then we used the scvelo function “scvelo.pp.filter_and_normalize”, with parameters “min_counts”=20 and “min_counts_u”=10, to filter genes on the basis on their number of spliced and unspliced counts, before normalizing the raw count matrix and selecting the top 3,000 HVGs.

We used the function “scvelo.pp.moments” to compute neighbouring cells and spliced and unspliced moments. After this, the inference of the genes’ splicing dynamics and the computation of the RNA velocities were computed from the dynamical model (using the functions “scvelo.tl.recover_dynamics”; “scvelo.tl.velocity” with “dynamical” mode; “scvelo.tl.velocity_graph”). Finally, we projected the velocities onto the first two DCs of the previously computed diffusion map (using the function “scvelo.tl.velocity_embedding”) and we computed the PAGA velocity graph using the function “scvelo.tl.paga” with default parameters.

### Sub-clustering of Visceral Endoderm cell populations

We performed a sub-clustering analysis of the AVE and emVE cells in a previously published scRNA-seq dataset (Nowotschin et al., 2019) obtained through the 10x protocol. We downloaded the raw count matrix as an AnnData object from https://endoderm-explorer.com and the metadata provided by the authors at: https://data.humancellatlas.org/explore/projects/4e6f083b-5b9a-4393-9890-2a83da8188f1. We selected only VE cells at E5.5 and E6.5, which were collected in three and two batches, respectively. We normalized the data separately for each stage and batch using the R package “scran”, and we log-transformed the matrices as previously described. At each stage, we integrated the data from the batches using the “mnn_correct” function from the “mnnpy” package (see above).

In Nowotschin et al. (2019), cells from the VE were split into two clusters: the extra-embryonic VE (which corresponds to our exVE clusters) and another including cells from the embryonic VE, which includes the AVE. Hence, our first step consisted in the identification of a cluster of AVE cells at both the E5.5 and E6.5 stages.

To this aim, we selected only embryonic VE cells and we found the intersection of the top 3,000 HVGs from our Smart-seq2 dataset (using only cells from our AVE and emVE clusters) at the corresponding stage with the genes quantified in the 10x data (Supplementary Figure 6A–B–C). This set of genes was used for all the analyses described below.

We computed the knn graph with Euclidean distance on the first 10 principal components with the default value for *k*, and we used the Leiden algorithm (Traag et al., 2019) to cluster the cells (Scanpy function “sc.tl.leiden”). The resolution was fixed in such a way to obtain two clusters at each stage, which were annotated as AVE and emVE based on the expression of known markers.

As a next step, we searched for sub-populations of cells in the AVE and the exVE clusters at E6.5 using the procedure explained below in detail. For each of these clusters, we performed Leiden clustering on the knn graph based on the Euclidean distance between the first 10 principal components computed on the top 2,000 HVGs. To estimate the number of clusters, we tried several combinations of the clustering parameters, i.e., the number of nearest neighbours k and the resolution *r*: we took *k* ∈ [15,35], with a step of 5, and *r* ∈[0.1,1], with a step of 0.1. For each pair of (*k*,*r*) values, the robustness of clusters was tested with a gene sub-sampling procedure, with the same strategy described above. To mitigate the risk of overfitting, we kept only clusters that have a number of specific marker genes greater than 1; the clusters with 0 or 1 specific markers were merged with the closest of the remaining clusters, based on the relative Euclidean distances between the cluster centroids. We considered a marker gene as “specific” if (i) it has a log-normalized mean expression larger than 0.01 and (ii) it is statistically significantly upregulated in all the pairwise comparisons with the other clusters.

By doing so, we obtained three sub-clusters in the AVE and two sub-clusters in the exVE at E6.5, which were annotated on the basis of HCR experiments, as shown in Figure 5.

The same type of sub-clustering gave only a single cluster in E5.5 AVE cells. Hence, we used a more supervised approach to verify whether the gene signature distinguishing AVE-medial and -lateral subclusters at E6.25 can separate AVE sub-populations at E5.5. To this aim, we selected the top 10 upregulated and the top 10 downregulated genes between late-AVE-lateral and -medial cells at E6.25 (Wilcoxon test with Benjamini-Hochberg correction method). Using these genes, we could identify two clusters in the AVE at E5.5 with the Leiden algorithm (using *k*=15) (*see* Figure 5B).

### Cell-cell communication analysis

The analysis of intercellular communication in the scRNA-seq data at E5.5 and E6.25 has been performed using a curated list of 2,548 ligand-receptor pairs (LRPs) (Caruso et al., 2020), to which we added the following LRPs based on previously reported direct interactions: CER1-MRC2 (Vento-Tormo et al., 2018), CER1-BMP2/BMP4 (Chi et al., 2011), CER1-NODAL (Aykul and Martinez-Hackert, 2016), LEFTY1-NODAL (Chen and Shen, 2004), LEFTY1-BMPR2 (Zabala et al., 2020). We selected only AVE, emVE and Epi cells, and we filtered out genes detected in fewer than 10 cells or with log-normalized mean expression computed on non-zero values smaller than 1.

Following the approach of Solovey and Scialdone (2020), we represented each LRP *α* as a weighted directed graph, where the number of nodes *N* is equal to the number of cell types (i.e., in our case AVE, emVE and Epi). The elements of the adjacency matrix of the graph for the LRP *α* are given by

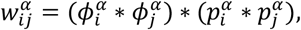

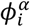 is the fraction of cells in the node *i* expressing the ligand and 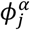 the fraction of cells in the node *j* expressing the receptor. All values of 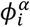 and 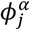 smaller than 0.1 were set to 0. 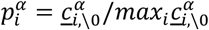, i.e., the ratio between the average of the non-zero log-expression values of the ligand in node *i* and the maximum of these averages computed over the different nodes.

If the LRP includes a complex, 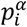 and 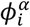 are the product of the *p* and *ϕ* of the genes belonging to the complex, respectively.

We filtered (i.e., set to 0) weights smaller than the 50th percentile of the weight distribution computed on all LRPs, and we eliminated LRPs with a null weight matrix. We ended up with 413 and 471 LRPs at E5.5 and E6.25, respectively.

Next, we considered LRPs containing at least one of the genes upregulated in the AVE at both E5.5 and E6.25, as identified from the diffusion pseudotime analysis. This additional filtering resulted in 43 LRPs containing ‘high-in-AVE’ ligands or receptors (*see* Figure 6A). We used COMUNET (Solovey and Scialdone, 2020) to visualise the communication patterns, in particular those involving the high-in-AVE Ephrins and Eph receptors (*Efna5* and *Ephb3*) (Supplementary Figure 7C–D).

### Analysis of genes in the Eph/Ephrin-signalling pathway

We considered genes in the Eph/Ephrin-signalling pathway and with log-normalized mean expression larger than 0.5, obtaining 14 genes at E5.5 and 13 genes at E6.25. Using these genes, we performed a Principal Components Analysis at each stage, using the Scanpy function “scanpy.tl.pca”, as previously described (Supplementary Figure 7A–C). By doing so, we noticed that the five cell types in the dataset are separated at E6.25, while the result at E5.5 is less clear.

Next, we asked which genes contributed significantly to the first three PCs at each stage. To this end, we generated 1,000 bootstrap samples of the cells and we performed a PCA on each of them. We assigned a p-value to each PC loading by computing the fraction of sign inversions obtained in the bootstrap procedure and accounting for possible axis reflections, as described in Peres-Neto et al. (2003). We considered genes as significant if they have at least one loading (considering the first three PCs) with p-value smaller than 0.01. We obtained 6 significant genes at E5.5 and 8 at E6.25. The expression of the significant Ephrin genes at each stage is shown in the dot plots in Figure 6B.

### In situ Hybridization Chain Reaction (HCR)

All the HCR probes were synthesised by Molecular Instruments (molecularinstruments.org, Pasadena, CA). *In situ* HCR v3.0 was carried out as previously described Choi et al. (2018), with the following modifications: embryos were dissected in ice-cold M2 medium, before directly fixing in 4% PFA (SantaCruz, sc281692) overnight at 4°C; proteinase-treatment was done with 10 μg/mL proteinase K (Thermo Scientific, EO0491), for 90 seconds at RT but using solutions pre-warmed to 37°C; post-fixation in 4% PFA was performed at 4°C for 20 minutes. When processing large number of embryos together, extra care was taken to make sure the temperature of the hybridisation and probe wash buffers the embryos were in didn’t drop below 37°C between transfers, by using prewarmed heat blocks. Methanol-dehydration/rehydration steps were omitted for samples requiring phalloidin-staining for F-actin visualisation. Following the HCR protocol, the samples were cleared in 87% (v/v) glycerol (Fisher Scientific, G/0650/17) in PBS (Sigma-Aldrich, P4417) for at least 3 days at 4°C before mounting for imaging using the same media.

### Wholemount immunofluorescence

Embryos were fixed in 4% PFA in PBS at RT for 30 minutes (or 5 minutes in ice-cold 1:1 acetone:methanol solution for Cytokeratin and Drebrin-1 staining), washed three times for 10 minutes in 0.1% Triton X-100 (Sigma-Aldrich, T8787) in PBS at RT; incubated in 0.25% Triton X-100 in PBS for 15 min at RT for permeabilization; washed twice for 5 minutes in 0.1% Tween-20 (Sigma-Aldrich, P1379) in PBS (PBST) at RT; blocked overnight in blocking reagent (5% donkey serum (Sigma-Aldrich, D9663), 3% bovine serum albumin (Sigma-Aldrich, A7906) in PBST) at 4°C; incubated overnight in primary antibodies diluted in the blocking reagent; washed three times for 10 minutes each, in PBST at RT; incubated overnight at 4°C in secondary antibodies and fluorophore-conjugated phalloidin diluted in PBST; washed thrice for 5 minutes in PBST at RT and the samples were cleared in depression slides with VECTASHIELD anti-fade mounting media containing DAPI (Vector Labs, H-1200) for 3 days at 4°C, before mounting for imaging using the same media.

The primary antibodies used were: 1:50 Rabbit anti-CK19 (Proteintech, 10712-1-AP), 1:100 Rabbit anti-Drebrin1 (Proteintech, 10260-1-AP), 1:100 Goat anti-PlexinA1 (R&D Systems, AF4309), 1:200 Rabbit anti-Oct4 (Abcam, ab19657). The secondary antibodies used were: Alexa Flour (AF)-555 Donkey anti-rabbit IgG (Invitrogen, A31572) and AF-633 Donkey anti-goat IgG, AF-488 (Invitrogen, A21082) both at 1:200. For F-actin staining either Phalloidin-Atto 550 (Sigma-Aldrich, 19083) or Phalloidin-Atto 647N (Sigma-Aldrich, 65906) were used at a final concentration of 1 mg/mL.

### Embryo culture and inhibitor studies

Embryo culture media made up of 49.5% CMRL-1066 (Pan BioTech, P04-84600), 49.5% KnockOut serum-replacement (Gibco, 10828-010), supplemented with was 1% L-glutamine was pre-equilibrated at 37°C and 5% CO_2_ for at least 2 hours prior to use. *Hhex-GFP* transgenic embryos dissected at E5.5 were briefly screened on a 35 mm glass bottom dish (MatTek Life Sciences, P35G-1.5-14-C) using low laser intensities under a confocal microscope to select those embryos where the GFP-positive AVE cells were still at the distal tip. The ectoplacental cone was left on the embryos to facilitate survival during culture. The embryos were cultured in Nunc™ Lab-Tek™ II chambered coverglass slides (Thermo Fisher, 155409) with 400 *μ*l of embryo culture media per chamber, under control- and treated-conditions at 37°C and 5% CO_2_. Culture periods from 4 to 15 hours were tested, but a duration of 4 hours was sufficient to document the complete migration of the AVE from the distal tip to the epiblast–ExE boundary and laterally across the embryo. To perturb Eph/Ephrin-signalling, the small molecule inhibitor NVP-BHG712 (Selleck Chemicals, A8683) was used at a final concentration of 100*μ*M and an equal volume of DMSO was used in the controls. To block Semaphorin-signalling, Mouse anti-Semaphorin6D (Santa Cruz Biotechnology, sc393258) or Goat anti-PlexinA1 (R&D Systems, AF4309) were used at a final concentration of 10 *μ*g/mL.

### Image acquisition and processing

Fixed-samples were imaged on a Zeiss LSM 880 confocal microscope using 20x (0.75NA) or 40x oil (1.36NA), objectives as appropriate. Z-stacks of embryos were acquired at 1 *μ*m interval using non-saturating scan parameters. Opacity rendering as 3D volumes and movies were made using the Volocity Software (Improvisions). For HCR images, the extended focus projections were maximum intensity projections. Figures were prepared with Adobe Photoshop 2020 and Adobe Illustrator 2020 (Adobe Inc.). Individual 3D opacity renderings for each channel were combined from individual layers for merged images.

## ACKNOWLEDGEMENTS

We would like to thank Ed Wilkes, Vinothini Rajeeve and Pedro Cutillas (Queen Mary University London) for help with the proteomics experiments; the FACS facility at the Oxford Weatherall Institute of Molecular Medicine; Richard Tyser for advice and help with cell collection for scRNA-Seq; and Matthew Stower for valuable discussions and helpful comments on the manuscript. JF was supported on a Joachim Herz Stiftung Add-on Fellowship for Interdisciplinary Life Science. This work was funded by the BBSRC Research Grant BB/I007806/1 to M.W. and B.V., Helmholtz Association to A.S., and Wellcome Strategic Awards 105031/C/14/Z, 108438/Z/15/Z and Senior Investigator Award 103788/Z/14/Z to S.S.

## LIST OF SUPPLEMENTARY TABLES

Supplementary Table 1: Top 50 marker genes per cluster at E5.5 Supplementary Table 2: Top 50 marker genes per cluster at E6.25

Supplementary Table 3: Lists of genes differentially expressed in diffusion pseudotime analysis of early-AVE and emVE cells at E5.5

Supplementary Table 4: Lists of genes differentially expressed in diffusion pseudotime analysis of late-AVE and emVE cells at E6.25

Supplementary Table 5: Lists of differentially expressed proteins between ExE and EPI halves in the proteomics dataset - Upregulated in ExE half

Supplementary Table 6: Lists of differentially expressed proteins between ExE and EPI halves in the proteomics dataset - Upregulated in EPI half

Supplementary Table 7: Lists of differentially expressed proteins between ExE and EPI halves in the phosphoproteomics dataset - Upregulated in ExE half

Supplementary Table 8: Lists of differentially expressed proteins between ExE and EPI halves in the phosphoproteomics dataset - Upregulated in EPI half

Supplementary Table 9: Lists of specific marker genes of the late-AVE sub-clusters at E6.5

Supplementary Table 10: Lists of differentially expressed genes between the late-AVE transitional sub-cluster and the emVE cluster at E6.5

Supplementary Table 11: Lists of differentially expressed genes between the late-AVE-lateral and late-AVE-medial sub-clusters at E6.5

Supplementary Table 12: Lists of differentially expressed genes between exVE-distal and exVE-proximal sub-clusters at E6.5

Supplementary Table 13: Raw data for the proteomics experiments at E6.5 Supplementary Table 14: Raw data for the phosphoproteomics experiment at E6.5

---

**Supplementary Figure S1.**
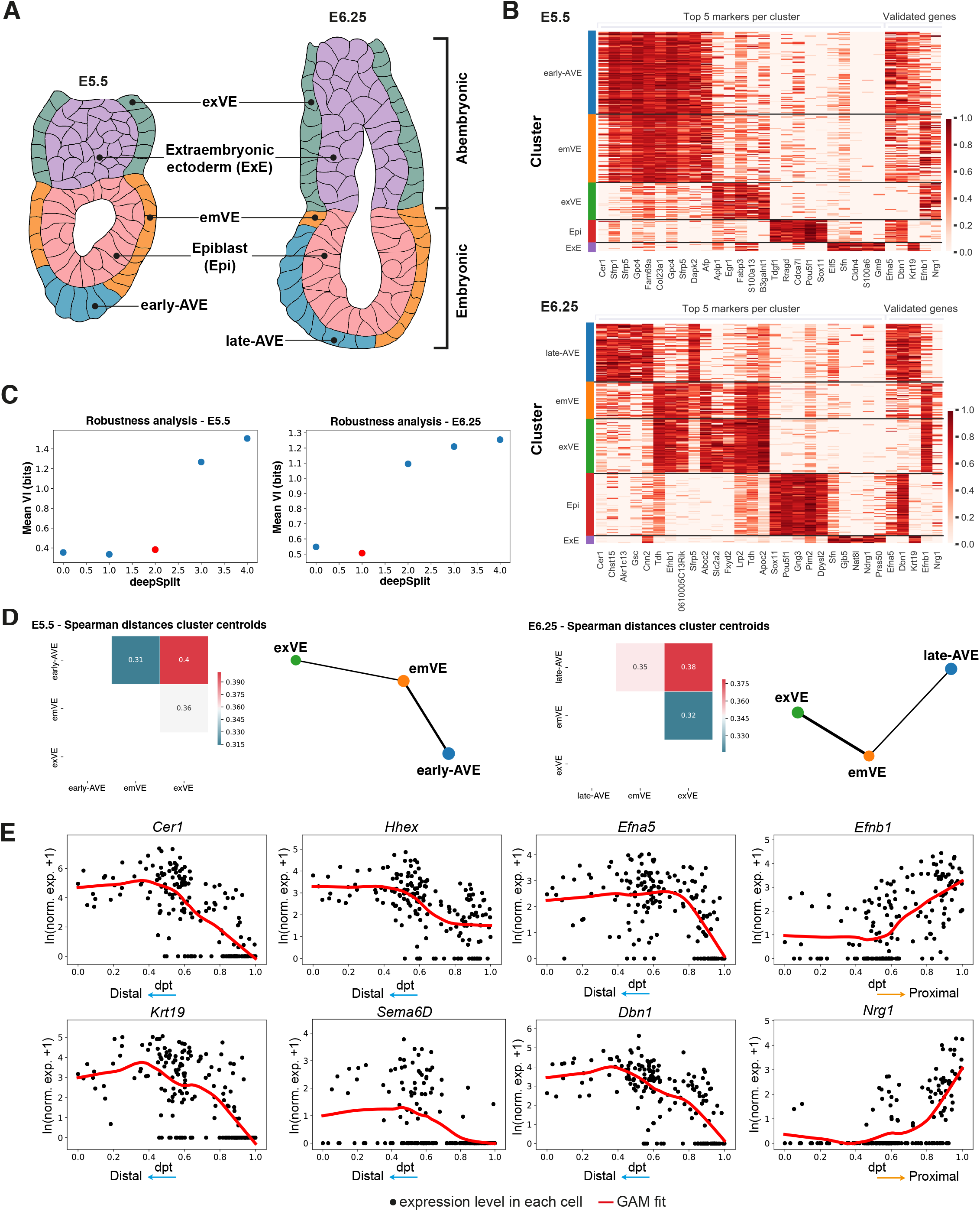
Expression of newly computed marker genes, robustness analysis for cell clustering, similarity for visceral endoderm clusters and expression in pseudotime of selected ‘high-in-AVE’ genes. (A) Illustrations of mid-sagittal sections through E5.5 and E6.25 embryos outlining the nomenclature used in this study to describe the different cell-types of the embryo at these stages. (B) Heatmaps showing the expression of the top 5 marker genes per cluster that we computed, at E5.5 (top) and E6.25 (bottom), and of the genes that have been selected for experimental validation (*Efna5*, *Dbn1*, *Krt19*, *Efnb1*, *Nrg1*). The normalized log expression levels of each gene are standardised so that they vary within the interval [0,1]. (C) Plot of the mean Variation of Information (VI) as a function of the deepSplit parameter, used for choosing the number of clusters in the clustering of the scRNA-seq data at E5.5 (left) and E6.25 (right). The red dot indicates the value of deepSplit chosen at each stage (see Methods for the details regarding the robustness analysis for cell clustering). (D) Matrix plots of the Spearman distances between the centroids of the AVE, emVE and exVE clusters, at E5.5 (left) and E6.25 (right), and partition-based graph abstraction (PAGA) plots for the AVE, emVE and exVE clusters, at E5.5 (left) and E6.25 (right). (E) Expression patterns of selected genes from the ‘high-in-AVE’ group (*Cer1*, *Hhex*, *Efna5*, *Krt19*, *Sema6d*, *Dbn1*) and low-in-AVE group (*Efnb1, Nrg1*) at E5.5 in the early-AVE and emVE clusters, as a function of diffusion pseudotime.

**Supplementary Figure S2.**
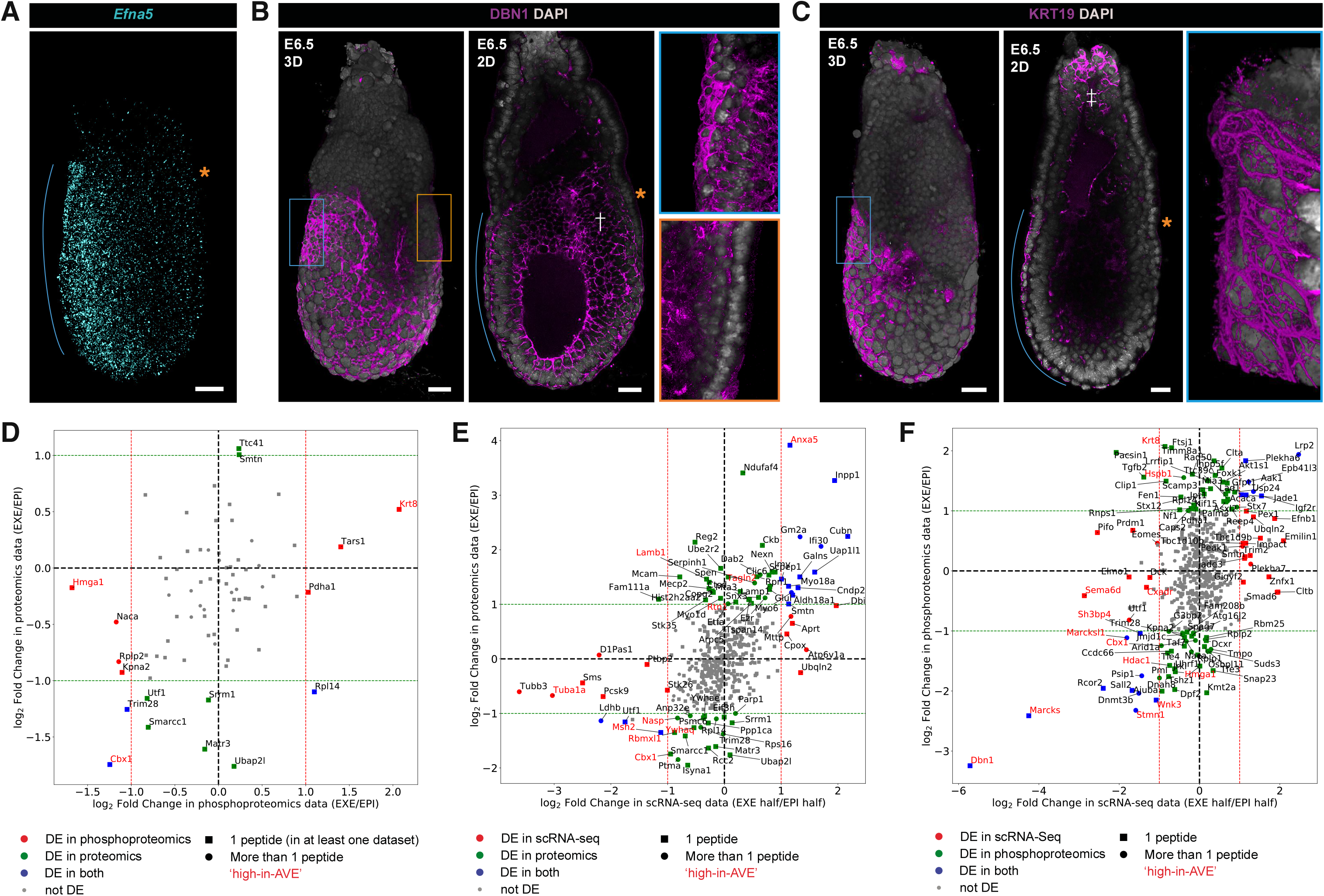
Analysis of proteomics and phosphoproteomics from mouse embryos at E6.5 and comparison with scRNA-seq data. (A-C) Continued anterior-enriched expression of high-in-AVE markers *Efna5* (visualised by *in situ* HCR), DBN1 and KRT19 (visualised by immunofluorescence) at E6.5. The blue lines indicate the position of the AVE and the orange asterisks mark the posterior side of the embryos. † and ‡ mark the expression of DBN1 and KRT19 in the epiblast and ExE respectively. Scale bars represent 20 μm and all embryos are orientated with the anterior on the left (←). (D) Scatter plot of log2 Fold Change, computed from a differential expression analysis between the embryonic and the abembryonic halves (EPI and EXE halves), in proteomics vs phosphoproteomics data (see the Methods section). Each marker represents a protein, the different colours indicate whether the protein is differentially expressed only in the phosphoproteomics (red), only in the proteomics (green) or in both datasets (blue), the different markers indicate whether a protein has only one peptide in at least one dataset (square) or if it has more than one peptide (circle). The names of proteins corresponding to genes from the high-in-AVE group are highlighted in red. Pearson’s correlation coefficient=0.36; p-value=1.3 × 10^−3^. (E) Scatter plot of log2 Fold Change, computed from a differential expression analysis between the EXE and the EPI halves, in proteomics vs scRNA-seq data at E6.5 and E6.25, respectively (see the Methods section). Each marker represents a protein/gene, the different colours indicate whether the protein is differentially expressed only in the scRNA-seq (red), only in the proteomics (green) or in both datasets (blue), the different markers indicate whether a protein has only one peptide (square) or if it has more than one peptide (circle). The names of proteins corresponding to genes from the high-in-AVE group are highlighted in red. Pearson’s correlation coefficient=0.52; p-value=8.5 × 10^−33^. (F) Scatter plot of log2 Fold Change, computed from a differential expression analysis between the EXE and the EPI halves, in phosphoproteomics vs scRNA-seq data at E6.5 and E6.25, respectively (see the Methods section). Each marker represents a protein/gene, the different colours indicate whether the protein is differentially expressed only in the scRNA-seq (red), only in the phosphoproteomics (green) or in both datasets (blue), the different markers indicate whether a protein has only one peptide (square) or if it has more than one peptide (circle). The names of proteins corresponding to genes from the high-in-AVE group are highlighted in red. Pearson’s correlation coefficient=0.39; p-value=1.0 × 10^−19^.

**Supplementary Figure S3.**
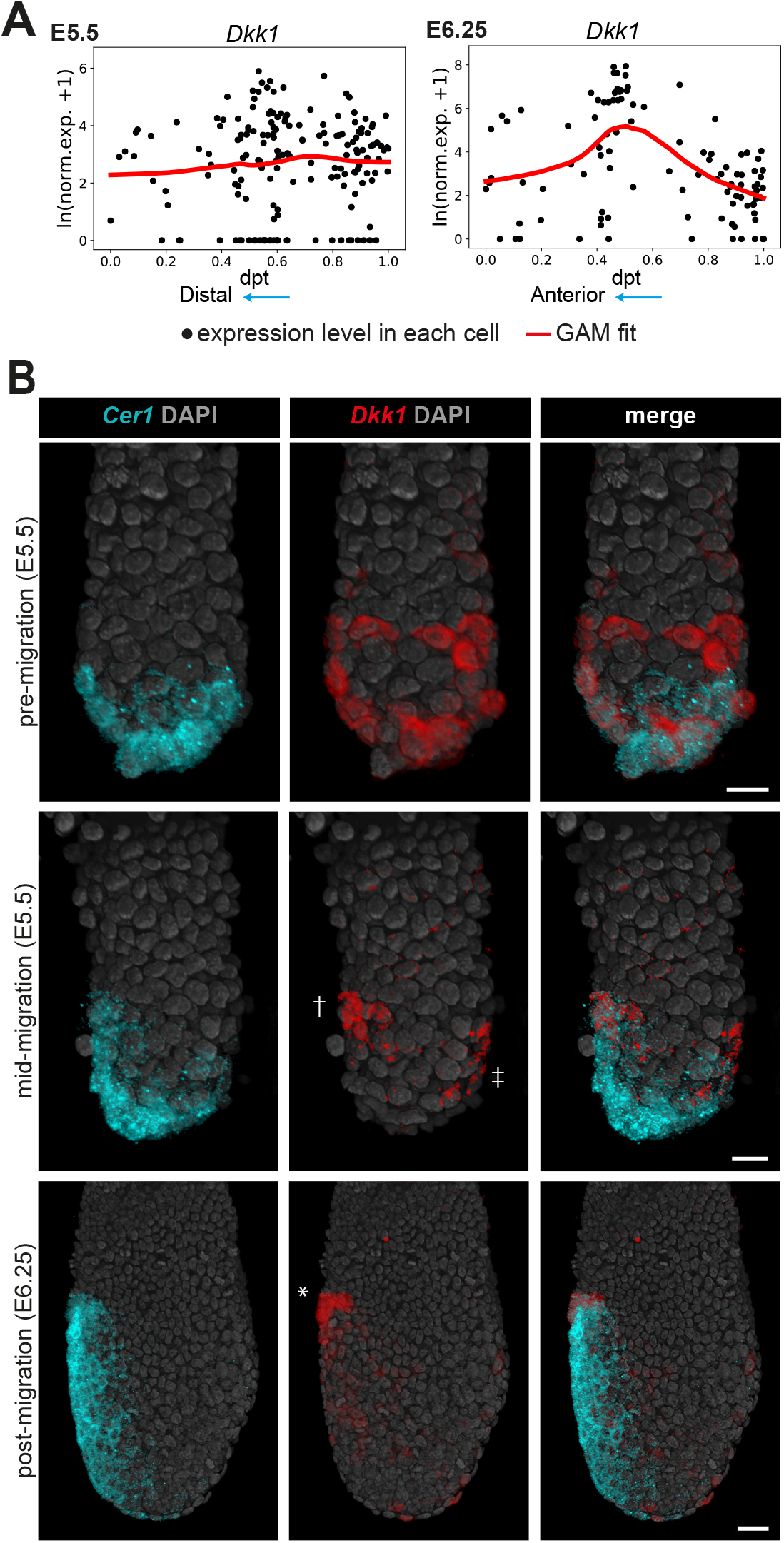
Symmetry breaking along the anterior–posterior axis relative to *Dkk1* expression in direct comparison to the migration of the AVE. (A) Expression pattern of *Dkk1* in the AVE and emVE clusters, as a function of diffusion pseudotime, at E5.5 (left) and E6.25 (right). (B) Volume renderings showing the changes to the *Dkk1* expression domain (visualised by *in situ* HCR) relative to the position of the *Cer1*-expressing AVE cells as they migrate from the distal portion of the embryo to the epiblast-ExE boundary. † and ‡ mark the segregated anterior and posterior expression domains of *Dkk1* at mid-migration stages before it gets confined to the leading (most-proximal) cells of the AVE (*) at E6.25. Scale bars represent 20 *μ*m and all embryos are orientated with the anterior on the left (←).

**Supplementary Figure S4.**
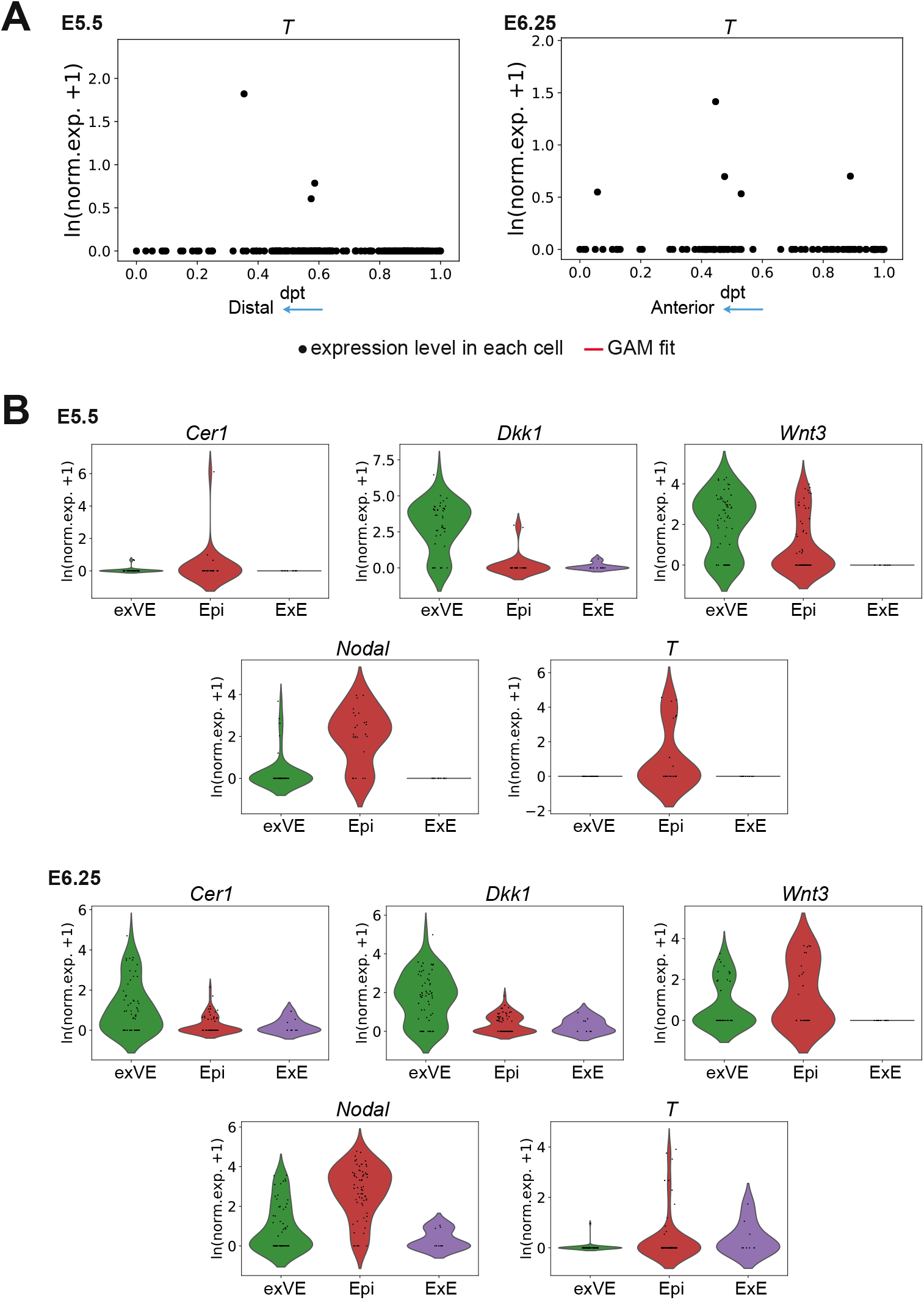
Expression of key anterior–posterior symmetry breaking markers in different cell populations of the embryo. (A) Expression patterns of *T* in the AVE and emVE clusters, as a function of diffusion pseudotime, at E5.5 (left) and E6.25 (right) (B) Violin plots showing the normalized log expression levels of *Cer1*, *Dkk1*, *Wnt3*, *Nodal* and *T* in the exVE, Epi and ExE clusters, at E5.5 (top) and E6.25 (bottom).

**Supplementary Figure S5.**
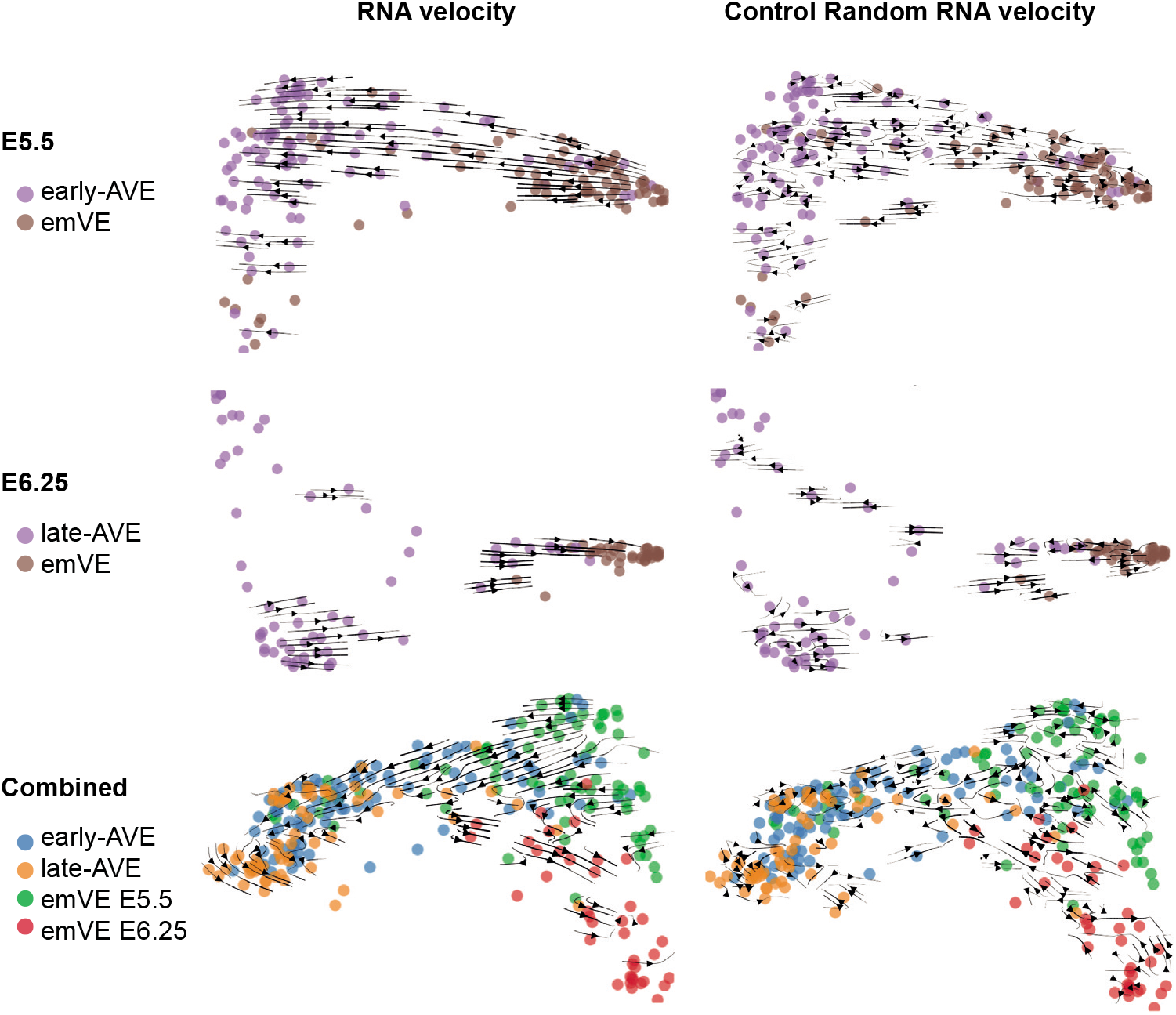
RNA velocity for E5.5 and E6.25 stages, separated and combined, and random control. RNA velocity streams projected on the first two diffusion components of a diffusion map computed on AVE and emVE cells at E5.5 (top left), E6.25 (centre left) and on the combined data from the two stages (bottom left). The right column shows the projection on the same embeddings of randomized RNA velocities, as a control.

**Supplementary Figure S6.**
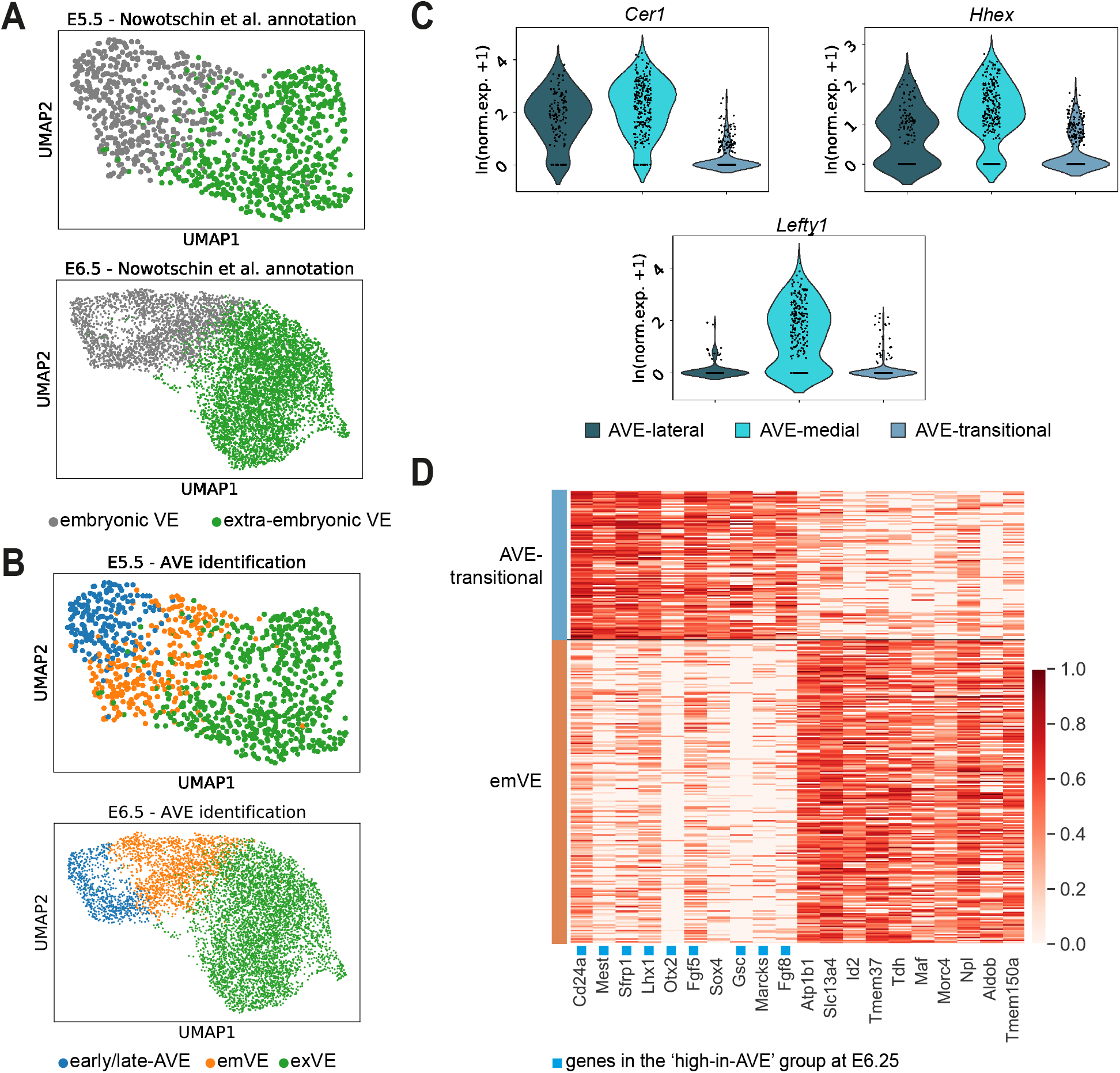
AVE identification from a public scRNA-seq dataset and characterization of the AVE-transitional cell population. (A) UMAP plots of the cells annotated as visceral endoderm in Nowotschin et al. (2019) at E5.5 (top) and E6.5 (bottom). Cells are coloured according to the original annotation as embryonic VE and extra-embryonic VE. (B) Same UMAP plots as in panel (A), but with the cells coloured according to our annotation (early/late-AVE, emVE and exVE clusters), obtained as described in Figure 5A and in the Methods. (C) Violin plots showing the normalized log expression levels of *Cer1*, *Hhex* and *Lefty1* in the late-AVE sub-clusters (AVE-lateral, AVE-medial and AVE-transitional) at E6.5. (D) Heatmap showing the expression of the top 10 genes upregulated in the AVE-transitional sub-cluster and the top 10 genes upregulated in the emVE cluster, obtained through a differential expression analysis between the AVE-transitional and the emVE clusters at E6.5. The normalized log expression levels of each gene are standardised so that they vary within the interval [0,1]. Genes that also belong to the ‘high-in-AVE’ group we identified at E6.25 are flagged in blue.

**Supplementary Figure S7.**
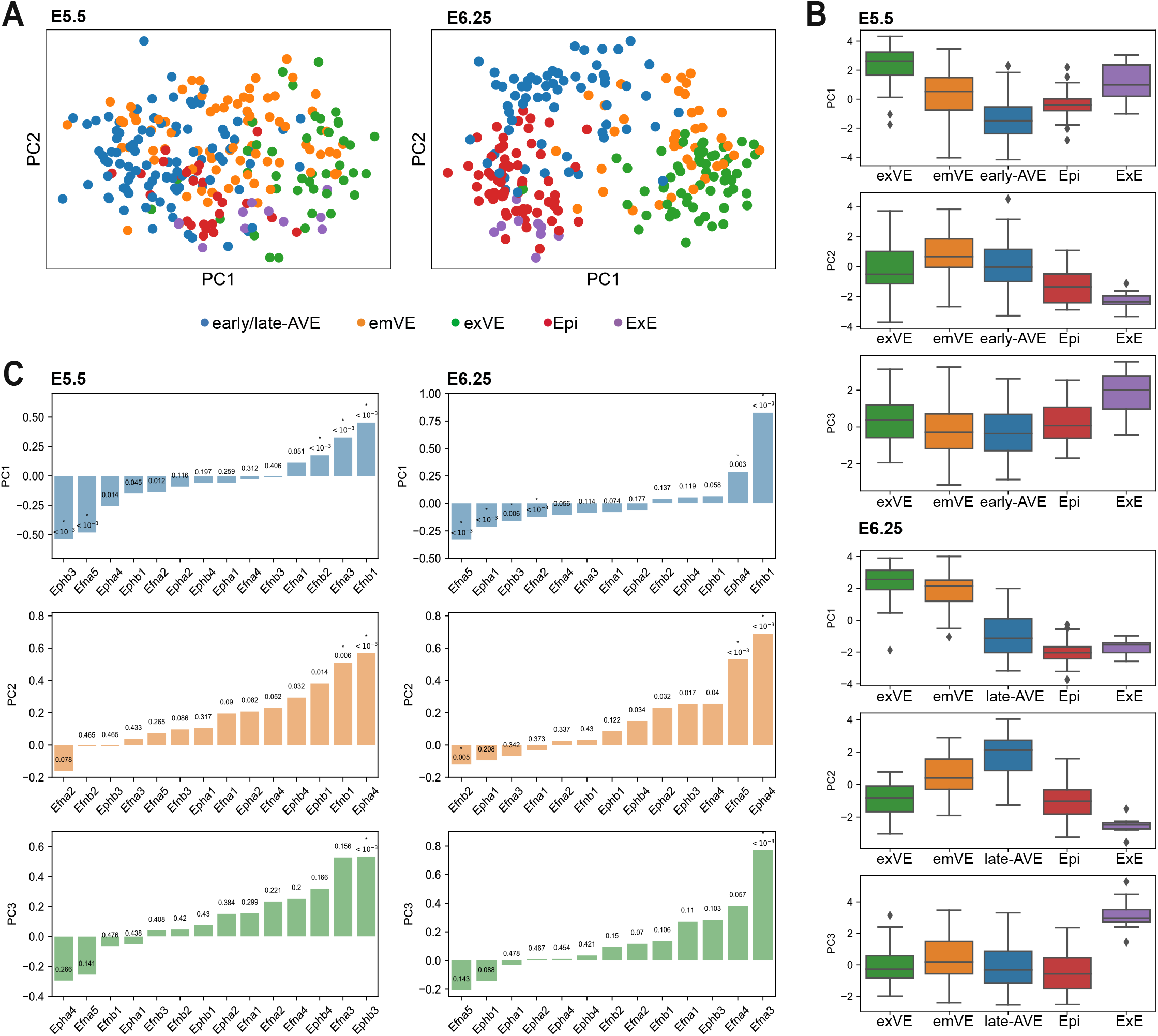
A Principal Components Analysis on genes belonging to the Eph/Ephrin-signalling pathway separates the five clusters present in the dataset. (A) Plots showing the first two principal components of a principal components analysis (PCA) performed on genes belonging to the Eph/Ephrin-signalling pathway, at E5.5 (left) and E6.25 (right). (B) Box plots of the values of the first three principal components grouped by cluster, at E5.5 (top) and E6.25 (bottom). (C) Loadings of the first three principal components for the genes used in the PCA, at E5.5 (left) and E6.25 (right). The p-values indicated for each gene were computed using a bootstrap procedure (see Methods). Significant genes (p-value < 0.01) are marked in the loading plots. We obtained 6 and 8 significant genes at E5.5 and E6.25, respectively.

**Supplementary Figure S8.**
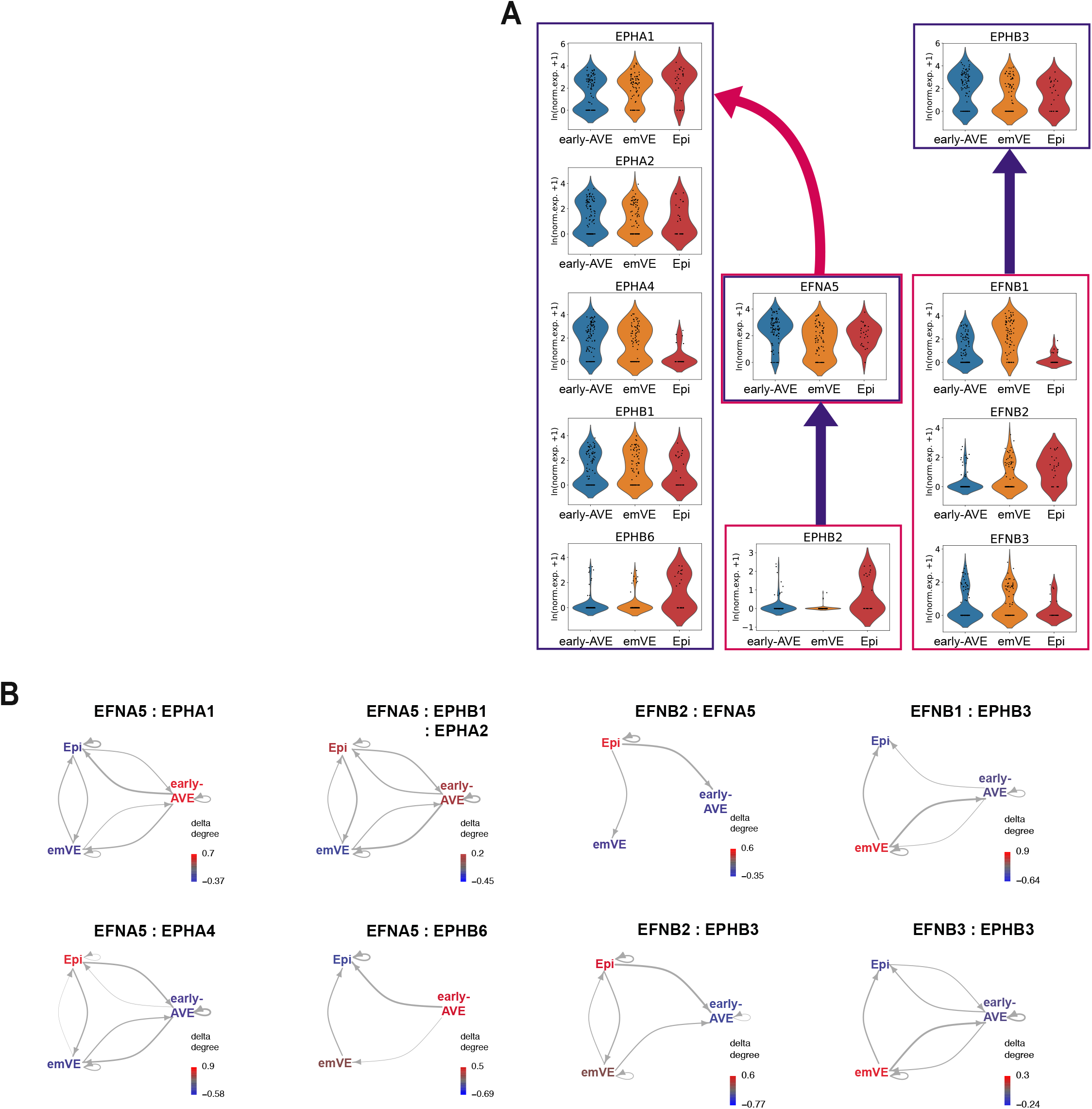
Expression levels and intercellular communication patterns for ligand–receptor pairs (LRPs) containing ‘high-in-AVE’ genes belonging to the Eph/Ephrin-signalling pathway. (A) Violin plots showing the normalized log expression levels of the genes belonging to the Eph/Ephrin-signalling pathway shown in Figure 6A, at E5.5. The red arrow shows that *Efna5* acts as a ligand for the receptors *Epha1*, *Epha2*, *Epha4*, *Ephb1*, and *Ephb6*, while the blue arrows show that *Efna5* is a receptor for *Ephb2* due to the bidirectional nature of signalling within this pathway. *Ephb3* is a receptor of the ligands *Efnb1*, *Efnb2*, *Efnb3*. (B) Intercellular communication patterns for the LRPs involving *Efna5* and *Ephb3* belonging to the ‘high-in-AVE’ gene group at E5.5.

**Supplementary Figure S9.**
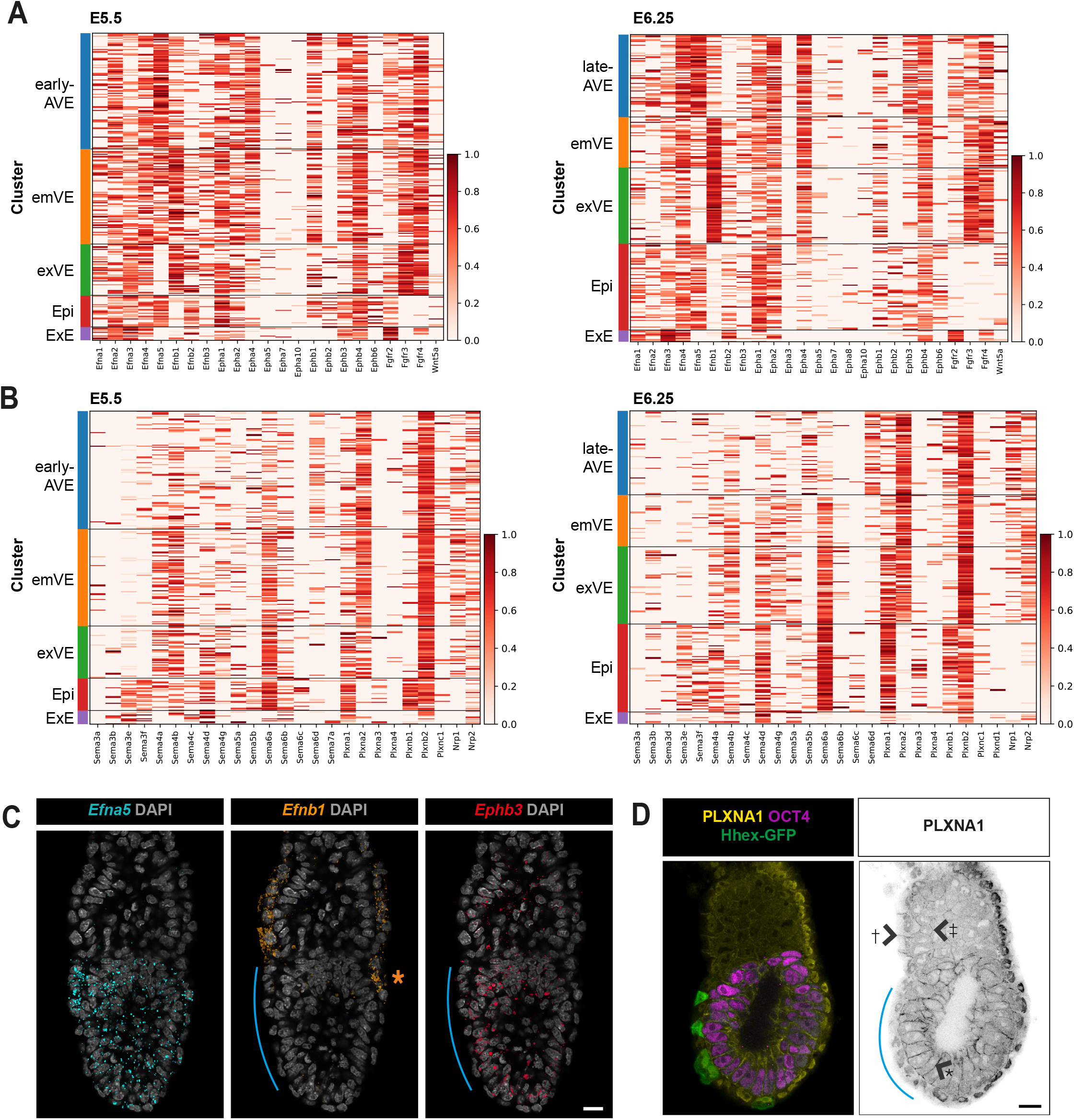
Expression of members of Ephrin- and Semaphorin-signalling pathways in the peri-implantation embryo. (A) Heatmap showing the expression of genes encoding components of the Eph/Ephrin-signalling pathway, at E5.5 (left) and E6.25 (right). The normalized log expression levels of each gene are standardised so that they vary within the interval [0,1]. (B) Heatmap showing the expression of genes encoding components of the Semaphorin/Plexin-signaling pathway, at E5.5 (left) and E6.25 (right). The normalized log expression levels of each gene are standardised so that they vary within the interval [0,1]. (C) Optical sections though late E5.5 embryos showing the expression of anterior-enriched *Efna5* and *Ephb3* relative to that of the posterior-enriched *Efnb1* (visualised with *in situ* HCR). The blue line indicates the extent of the AVE and the orange asterisk the posterior side of the embryo. (D) Optical section through a late E5.5 embryo showing the expression of PlexinA1 (PLXNA1- the receptor for the AVE-specific Semaphorin, SEMA6D), in the membranes of epiblast (*), exVE (†) and ExE (‡). Hhex-GFP and OCT4 mark AVE and epiblast cells respectively. The blue lines show the position of the AVE. All scale bars represent 20 *μ*m and all embryos are orientated with the anterior on the left (←).

